# Transcriptional, epigenetic and metabolic signatures in cardiometabolic syndrome defined by extreme phenotypes

**DOI:** 10.1101/2020.03.06.961805

**Authors:** Denis Seyres, Alessandra Cabassi, John J Lambourne, Frances Burden, Samantha Farrow, Harriet McKinney, Joana Batista, Carly Kempster, Maik Pietzner, Oliver Slingsby, Thong Huy Cao, Paulene A Quinn, Luca Stefanucci, Matthew C Sims, Karola Rehnstrom, Claire L Adams, Amy Frary, Bekir Ergüener, Roman Kreuzhuber, Gabriele Mocciaro, Simona D’Amore, Albert Koulman, Luigi Grassi, Julian L Griffin, Leong Loke Ng, Adrian Park, David B Savage, Claudia Langenberg, Christoph Bock, Kate Downes, Nicholas J Wareham, Michael Allison, Michele Vacca, Paul DW Kirk, Mattia Frontini

## Abstract

Improving the understanding of cardiometabolic syndrome pathophysiology and its relationship with thrombosis are ongoing healthcare challenges. Using plasma biomarkers analysis coupled with the transcriptional and epigenetic characterisation of cell types involved in thrombosis, obtained from two extreme phenotype groups (obese and lipodystrophy) and comparing these to lean individuals and blood donors, the present study identifies the molecular mechanisms at play, highlighting patterns of abnormal activation in innate immune phagocytic cells and shows that extreme phenotype groups could be distinguished from lean individuals, and from each other, across all data layers. The characterisation of the same obese group, six months after bariatric surgery shows the loss of the patterns of abnormal activation of innate immune cells previously observed. However, rather than reverting to the gene expression landscape of lean individuals, this occurs via the establishment of novel gene expression landscapes. Netosis and its control mechanisms emerge amongst the pathways that show an improvement after surgical intervention. Taken together, by integrating across data layers, the observed molecular and metabolic differences form a disease signature that is able to discriminate, amongst the blood donors, those individuals with a higher likelihood of having cardiometabolic syndrome, even when not presenting with the classic features.

## Introduction

Cardiovascular disease (CVD) is the primary cause of death worldwide (17.9 million deaths in 2016, 31% of all deaths)^1^ accompanied by an ever increasing number of overweight and obese individuals, which place a burden of hundreds of billions of dollars on healthcare systems each year^2, 3^. Cardiometabolic syndrome (CMS) increases both CVD and type 2 diabetes (T2D) risk^4^. CMS is a cluster of interrelated features including: obesity, dyslipidemia, hyperglycemia, hypertension and non-alcoholic fatty liver disease^5^. These features have overlapping components, including visceral fat deposition, high triglycerides, high low-density lipoprotein (LDL)-cholesterol, high fasting blood glucose, hypertension, decreased high-density lipoprotein (HDL)-cholesterol and low-grade chronic inflammation^6–8^. The therapeutic approaches aim to mitigate these features and include: weight loss strategies^9^, lipid lowering drugs^10^, antiplatelet therapies^11^, glucose lowering^12, 13^ and anti-inflammatory drugs^14^. The relationship between cardiometabolic health and body weight is complex^15^. CVD risk varies among individuals of similar body mass index (BMI) depending on adipose tissue (AT) distribution and functionality^16–20^. AT acts as an active endocrine organ^21, 22^ and when dysfunctional, plays a major role in metabolic disorders inducing peripheral insulin resistance, and contributing to low-grade chronic inflammation^23^.

Whilst the participation of platelets and neutrophils in thrombosis and that of macrophages in atherosclerotic plaque formation are well established^24–26^, the role of these cell types in atherogenesis and CVD onset has been appreciated only recently^27^. Additionally, prolonged exposure to low-grade inflammation is known to modify the functional phenotype of monocytes (an effect named trained immunity^28^), platelets^29, 30^ and neutrophils^31, 32^. The molecular characterisation of these phenotypic changes remains incomplete, motivating the need for extended molecular phenotyping of these cells performed here. Previous multi omics studies in blood cells have identified pathways involved in CVD and obesity, and confirmed whole blood as a source of surrogate biomarkers able to delineate the metabolic status^33^. Several risk score algorithms have been developed to predict the risk of complications associated with obesity^34–39^. However, a number of questions still remain open. CVD may also occur in the absence of other comorbidities and certain events have a better clinical outcome in overweight and obese patients compared with their leaner counterparts (the so-called "obesity paradox")^40^. We speculated that extreme phenotypes could be used to determine disease signatures, including new features, that are informative of disease aetiology in the general population.

Here, we present the molecular characterization of the transcriptional (RNA sequencing, RNA-Seq) and epigenetic (histone 3 lysine 27 acetylation, H3K27ac; reduced representation bisulfite sequencing, RRBS, and Illumina HumanMethylation450 BeadChip) changes in neutrophils, monocytes, macrophages and platelets in morbidly obese (BMI>40kg/m2; no obvious genetic cause^41^) and in familiar partial lipodystrophy type 2 (hereafter lipodystrophy; causal mutations in *PPARG* or *LMNA* genes, as verified by whole genome sequence^41, 42^) individuals. We also investigated the reversibility of these molecular changes in the obese group after bariatric surgery. We found that proinflammatory gene expression programs were downregulated, alongside more modest differences in regulatory elements usage and almost no differences in DNA methylation profiles. Altogether, the data indicate a reduced ability of these cells to be activated and undergo extracellular traps (netosis), which was further confirmed by neutrophil and platelet cell functional assays, which showed a reduced ability to adhere, the key initial step during their activation. Lastly, we indeed identified the molecular signatures for CMS and devised a penalised logistic regression approach to stratify individuals in the general population based on their CMS risk.

## Results

### Metabolic signatures in the obese and lipodystrophy groups

Participants were recruited as follow: controls (N=20; from which metabolically healthy individuals, hereafter lean, were selected, see METHODS), lipodystrophy (N=10), morbidly obese referred for bariatric surgery (N=11) and blood donors (hereafter BD; N=202)^43^. We collected age and body weight (BW), and performed plasma biochemistry assays for the following: leptin, adiponectin, insulin, free fatty acid (FFA), glucose (GLC), serum lipid (triglycerides (TG), total cholesterol (TC), high density lipoprotein (HDL-C), low-density lipoprotein (LDL-C)), activity of alanine and aspartate amino-transferases (ALT and AST, respectively) and high-sensitivity C-reactive Protein (hsCRP). Additionally, we computed the following: leptin-adiponectin ratio (LAR), Homeostatic Model Assessment for Insulin Resistance (HOMA-IR) and Adipose Tissue Insulin Resistance (AT-IR) indices (**Table 1** and **Table S1**).First, we wanted to determine if the different groups could be separated based on their plasma biochemistry and anthropometric characteristics. Compared to the other groups, the lipodystrophy group had elevated GLC, TC, TG, ALT, AST, insulin (and consequently HOMA-IR and AT-IR); whereas, HDL-C and LDL-C were decreased. Instead, the obese group, compared to the other 3, had elevated LAR, LDL-C and hsCRP.

**Table 1.** Descriptive characteristics of the study groups. Average value and standard deviation are indicated.

|  | Blood donors<br>(n=202) | Controls<br>(n=20) | Lipodystrophy<br>(n=10) | Obese<br>(n=11) | Post<br>surgery<br>(n=10) |
| --- | --- | --- | --- | --- | --- |
| Adiponectin<br>($\mu\text{g/ml}$) | 10.1 ± 6.3 | 10.7 ± 3.7 | 3.2 ± 2.3 | 5.9 ± 1.9 | 6.4 ± 2.6 |
| AGE(years) | 57.3 ± 11.1 | 40.7 ± 11 | 45.1 ± 9.6 | 46.3 ± 12.3 | 43 ± 12.6 |
| ALT (U/L) | 34.6 ± 12 | 27.1 ± 7.5 | 56 ± 12.7 | 35.7 ± 9.4 | 36.1 ± 17 |
| AST (U/L) | 25.5 ± 11.1 | 21.8 ± 6.9 | 39 ± 16.8 | 22.6 ± 3.8 | 18.9 ± 6.9 |
| AT-IR | 2.6 ± 2.5 | 1.9 ± 2.5 | 8.4 ± 7 | 7.2 ± 11.2 | 4.8 ± 5.4 |
| BMI ($\text{kg/m}^2$) | 26.4 ± 4.9 | <25 | <25 | 45 ± 5.1 | - |
| BW (kg) | 76 ± 14.9 | - | 73.2 ± 9.7 | 137.9 ± 35.2 | - |
| FFA ($\mu\text{mol/L}$) | 189.3 ± 132.8 | 156.5 ± 103.3 | 259.6 ± 174.9 | 293.1 ± 164.6 | 232.2<br>± 141.5 |
| GLC (mmol/L) | 5.4 ± 1.8 | 4.9 ± 1 | 8.3 ± 3.4 | 5.3 ± 0.6 | 5.3 ± 1.5 |
| HDL-C (mmol/L) | 1.6 ± 0.5 | 1.7 ± 0.4 | 0.8 ± 0.6 | 1.3 ± 0.2 | 1.3 ± 0.2 |
| HOMA-IR | 4.3 ± 4.9 | 2.5 ± 2.2 | 13 ± 10.5 | 7.1 ± 11.3 | 8.6 ± 18.4 |
| hsCRP (mg/L) | 1.9 ± 1.8 | 2.2 ± 1.2 | 2.3 ± 3.3 | 7.4 ± 6.9 | 2.9 ± 5.6 |
| Insulin (pmol/L) | 118.4 ± 117.1 | 76.4 ± 55.6 | 261.7 ± 262.3 | 190.6 ± 276.2 | 178.7<br>± 294.3 |
| LAR | 1.8 ± 2.1 | 2 ± 2.1 | 2.3 ± 1.9 | 13.7 ± 6.6 | 5.5 ± 4.5 |
| LDL-C (mmol/L) | 2.9 ± 0.9 | 2.7 ± 0.8 | 1.7 ± 0.5 | 2.4 ± 0.8 | 2.6 ± 1 |
| Leptin (ng/ml) | 14.2 ± 14.7 | 19.8 ± 17.1 | 7.6 ± 7.8 | 74.1 ± 30.4 | 29.9 ± 21 |
| TC (mmol/L) | 5.3 ± 1.1 | 4.9 ± 1 | 4.2 ± 1 | 4.5 ± 0.8 | 4.1 ± 1.4 |
| TG (mmol/L) | 1.6 ± 0.9 | 1.2 ± 0.9 | 5.6 ± 5.5 | 1.9 ± 0.7 | 1 ± 0.4 |

To visualize how these parameters separate obese, lipodystrophy and BD, we performed a principal component analysis (PCA), which showed that obese, lipodystrophy and BD groups were distributed over distinct, albeit partially overlapping, dimensions (**Fig.1A**). The first two components (PC1 and PC2) were sufficient to distinguish the different groups (Obese versus Lipodystrophy: p value = 0.002; Obese versus BD: p value < 2.2e-16; Lipodystrophy versus BD: p value < 2.2e-16; Hotelling’s T-squared test with F distribution). Lipodystrophy and obese were separated from BD along PC1, whilst they were separated from each other along PC2. Loading and contribution analysis (**Fig.1B**) showed that the main contributors to the separation along PC1 were BW, LAR, hsCRP, AST, ALT, GLC, AT-IR, HOMA-IR and TG.

**Figure 1.**
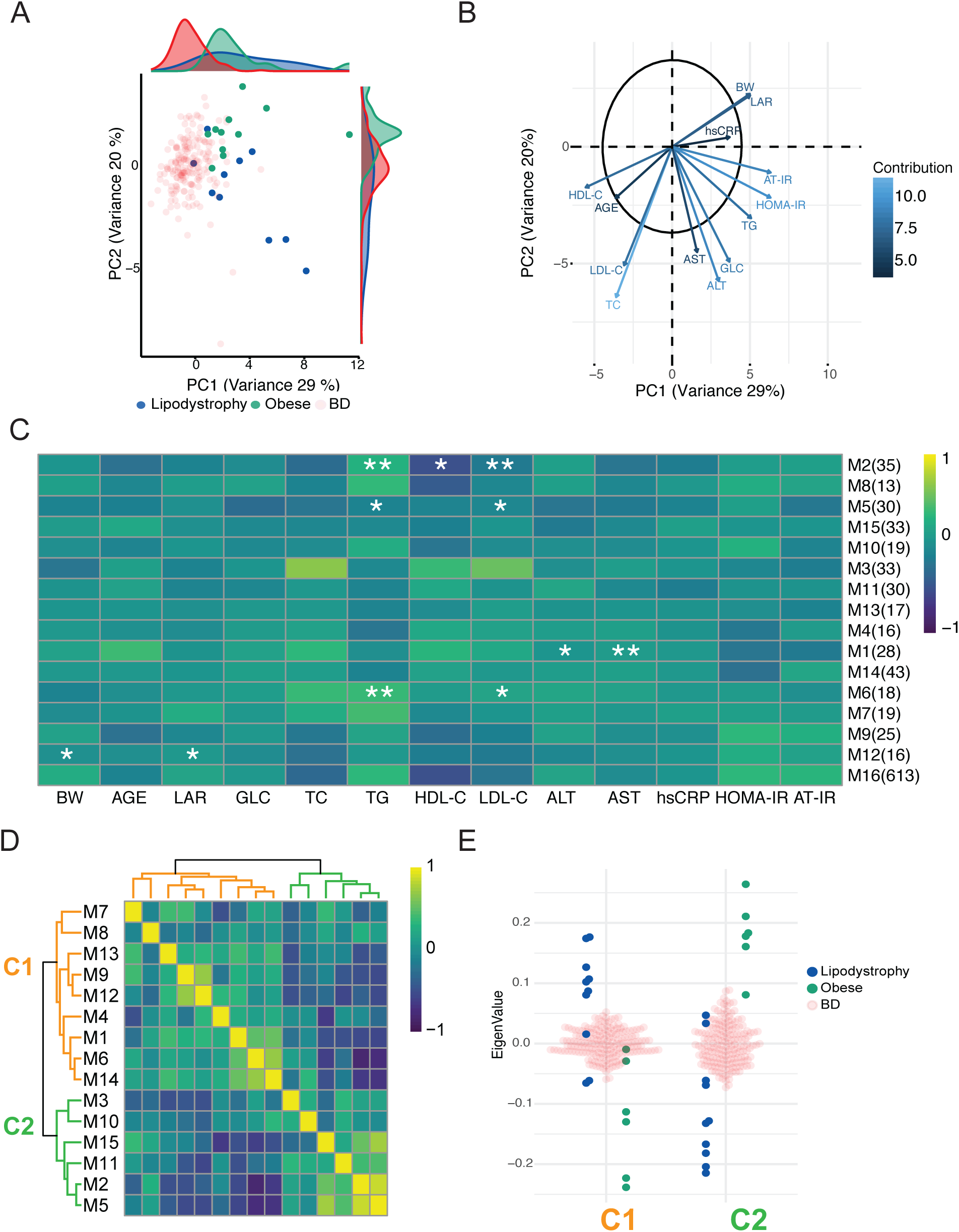
Metabolic signatures in the obese and lipodystrophy groups. A. Principal component analysis (PCA) of three groups: obese, green; lipodystrophy, blue; and blood donors (BD), light red. PCA was performed using the parameters below. B. Representation of PCA loadings on: age, weight (BW), body mass index (BMI), leptin-adiponectin ratio (LAR), glucose (GLC), triglycerides (TG), total cholesterol (TC), high density lipoprotein (HDL-C), low-density lipoprotein (LDL-C), alanine amino-transferase (ALT), aspartate amino-transferase (AST), Homeostatic Model Assessment for Insulin Resistance (HOMA-IR) and adipose tissue insulin resistance (AT-IR) indexes and high-sensitivity C-reactive Protein (hsCRP). Colour and arrow length scale represent contribution to variance on the first two principal components. C. Metabolite module-trait associations using WGCNA consensus analysis and 988 metabolites. Each row corresponds to a module eigen-metabolites (ME), and each column to a parameter. Number of metabolites in each module is indicated in brackets. Cell colour represents Pearson’s correlation as shown by legend. Significance is annotated as follows: * P≤ 0.05, ** P ≤ 0.01, *** P ≤ 0.001, **** P ≤ 0.0001 (Fisher’s test p value corrected for multi testing). D. Heatmap of extreme phenotype groups’ MEs adjacencies in the consensus MEs network. The heatmap is color-coded by adjacency, yellow indicating high adjacency (positive correlation) and blue low adjacency (negative correlation). E. Beeswarm plot using average MEs per cluster presented in D.

Additionally, BW, LAR, hsCRP separated the obese from the lipodystrophy groups in one direction along PC2, whilst AST, ALT, GLC, AT-IR, HOMA-IR and TG separated them in the opposite direction. The differences observed between the obese and the lipodystrophy groups in plasma biochemistry suggest that, while AT dysfunction is a shared feature, its influences were different in the two groups.

We further characterised the differences between the obese and lipodystrophy groups by investigating plasma metabolites, whose levels are known to be influenced by both extreme phenotypes^44, 45^; ^46, 47^. We identified and quantified 988 plasma metabolite species (using Metabolon^ⓡ^ (METHODS)) and we performed a weighted gene co-expression network consensus analysis (WGCNA)^48^ to create groups of metabolites whose levels were correlated across samples. To identify shared features and to reach the sample size as recommended for such analysis^48^ the obese and lipodystrophy group were analyzed together. This analysis identified 16 clusters of metabolites (named modules, M1 to M16; **Table S2**). To determine the relationship between modules, anthropometric traits and plasma biochemistry, we investigated if any correlation existed. Of the 208 tested associations, we found that 11 modules showed significant associations with BW, LAR, TG, HDL-C, LDL-C, ALT and AST in the extreme phenotype groups (FDR adjusted Fisher p values < 0.05; **Fig.1C**); while no associations were found in the BD cohort (not shown). To determine which modules were associated with each of the two extreme phenotype groups, we analysed the modules eigen-metabolite adjacencies (**Fig.1D**). The modules formed different clusters, C1 and C2 were found using extreme phenotype groups, C3 and C4 using BD samples (**Fig.S1A**). Plotting the average eigen-metabolite value for each cluster (**Fig.1E**) we showed that C1 and C2 represented the obese and lipodystrophy groups respectively, whereas clusters C3 and C4 could not discriminate between obese and lipodystrophy (**Fig.S1B**). C1 metabolites were significantly enriched in alanine, aspartate and glutamate metabolism, phenylalanine metabolism, nitrogen metabolism and TCA cycle; whereas C2 metabolites we found glycine, serine and threonine metabolism and cysteine and methionine metabolism pathways (**Table S2**).

Our analysis demonstrated that the two extreme phenotype groups could be identified by their metabolic signatures, associated with clinical parameters, which also set them apart from the general population represented by BD.

### Extreme phenotypes influence innate immune cell types and platelets transcriptional and epigenetic signatures

Next, we determined the influence of the changes in plasma on neutrophils, monocytes, macrophages and platelets, as these are some of the key players in atherogenesis and thrombus formation^49^ (**Fig.2A**). We compared gene expression (RNA-sequencing), active chromatin (histone 3 lysine 27 acetylation distribution) and DNA methylation (reduced representation bisulfite sequencing and Illumina arrays) in controls, BD and extreme phenotype groups. For each assays we performed the following comparisons: lean versus obese, lean versus lipodystrophy and obese versus lipodystrophy (**Fig.2A** and **Table S7**). For each comparison we identified differentially expressed genes (DEG; **Table S8-S11**), differentially acetylated regions (DAcR; **Table S12-S14**) and differentially methylated CpG islands (**Table S15-S17**) at a FDR 5%.

**Figure 2.**
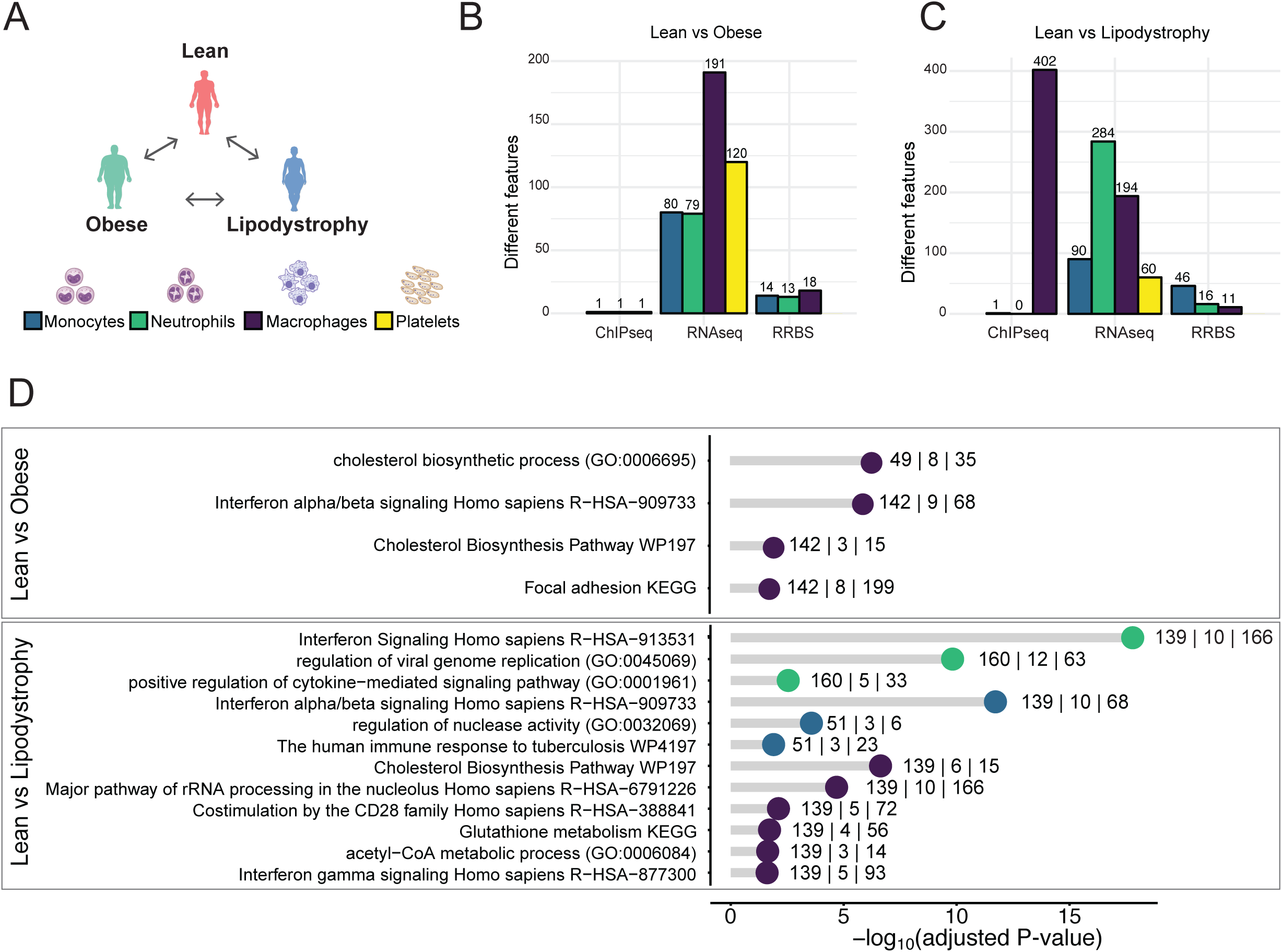
Transcriptional and epigenetic signatures in extreme phenotype groups for three innate immune cell types and platelets. A. Schematic overview of the comparisons made in the 4 different cell types (Monocytes: blue ; Neutrophils: green ; Macrophages: purple ; Platelets: yellow). B and C. Barplot showing the number of features significantly different: H3K27ac distribution (ChIP-Seq), gene expression (RNA-Seq) and DNA methylation (RRBS). Each bar is color coded to represent the different cell types as in A. B represents results when comparing lean and obese individuals. C represents results when comparing lean individuals and lipodystrophy patients. D. Functional GO term annotation of up-regulated genes when comparing lean versus obese group (top) and lean individuals versus lipodystrophy group (bottom), colour coded by cell types as above. The numbers near each dot indicate, from left to right: number of submitted genes, number of genes overlapping with the category and number of genes in the category.

Overall, we observed modest changes at transcriptional and epigenetic level in all comparisons (**Fig.2B**, **Fig.2C** and **Fig.S2B**), relatively to those observed in other tissues^50, 51^. The largest number of changes was found in active chromatin (3,616 DAcR) in the comparison between macrophages of the obese and lipodystrophy groups (**Fig.S2B**) and these were not accompanied by nearly as many changes in gene expression. This indicates that either similar transcriptional outputs were achieved using different regulatory landscapes^52^ or that these cells have been differently primed to respond to acute stimuli. These findings were in agreement with the absence of overlaps between DEG and genes previously associated with trained immunity^53^ in the lean versus obese and lean versus lipodystrophy comparisons.

Cell type specific functional annotation by gene ontology (GO) terms enrichment analysis for the DEG between the lean and obese groups (**Fig.2D**) found an enrichment for GO terms related to interferon alpha/beta signalling pathway, as well as focal adhesion in DEG upregulated in macrophages (**Table S18)**. In monocytes, up-regulated DEG were enriched for GO terms related to inflammatory response and down-regulated DEG were enriched in GO terms related to programmed cell death and ion homeostasis (**Table S19**). In neutrophils, down-regulated DEG were enriched for genes responding to antithrombotic drugs (**Table S20**). In the comparison between the lean and lipodystrophy groups (**Fig.2D**), macrophages up-regulated DEG were enriched in GO terms related to cholesterol biosynthesis and immune response activation. In monocytes and neutrophils, up-regulated DEG were enriched in terms related to interferon and immune responses. However modest, these changes illustrated how the exposures, to which the cell types involved in the development of atherosclerosis and in thrombus formation are subjected, modify the molecular phenotypes. Similar results have previously been reported for whole blood cell DNA methylation^54^. With the above exception in macrophages, we found that the two extreme phenotype groups were, as expected, more similar to each other than to the lean group overall, again reflecting the underlying AT dysfunction. To determine if these transcriptomic and epigenetic changes are reversible after exposures removal, a second blood sample was taken from the same obese individuals six months after bariatric surgery, and the same assays were performed.

### Effect of bariatric surgery on transcriptional and epigenetic landscapes, and cell functions

Bariatric surgery is effective in the management of extreme obesity and associated comorbidities, including CMS risk^55^, with well-established long-term benefits on weight loss, diabetes, hypertension and dyslipidemia^56^. While the effect of this intervention has already been reported^57, 58^, little is known about the underlying molecular mechanisms. Because we sampled the same individuals robust pairwise comparisons could be used. In plasma biochemistry we observed a decrease for LAR, TG, hsCRP, AT-IR and AST and an increase of HDL-C (p values: 7.22*10^-6^, 2.63*10^-9^, 4.98*10^-4^, 2.51*10^-2^, 1.48*10^-3^ and 1.86*10^-3^ respectively; conditional multiple logistic regression, adjusted for age and sex; **Fig.3A; Table S1**). Transcriptional and epigenetic paired analyses (**Fig.3B**) identified DEG in macrophages (599), monocytes (1,931), neutrophils (2,571) and platelets (2,883; **Table S8-S11**), DAcR in monocytes (229) and neutrophils (788; **Table S13-S14**) and differentially methylated CpGs in macrophages (201), monocytes (48) and neutrophils (198; **Table S15-S17**).

**Figure 3.**
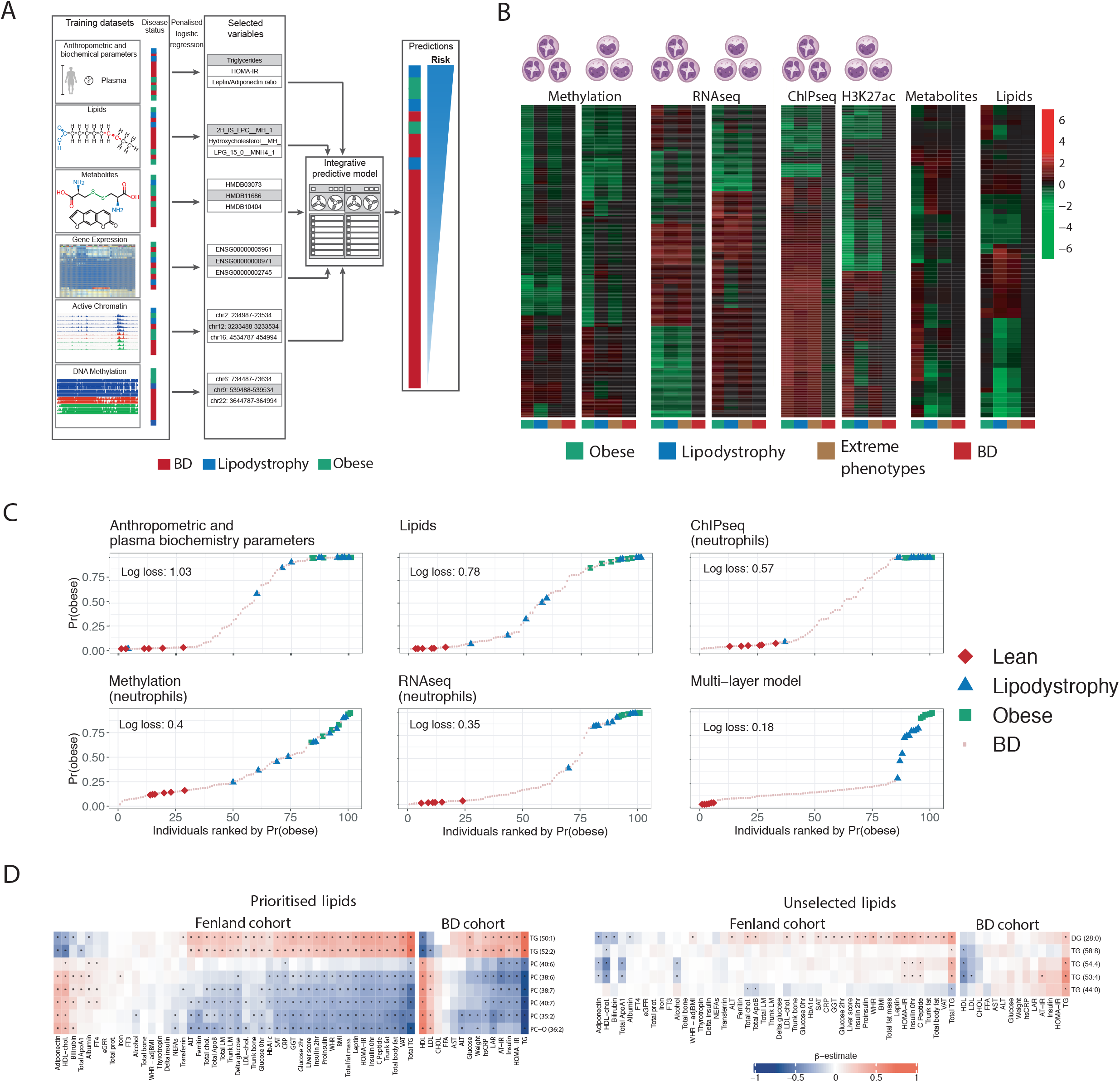
Multi-omic signature classification of extreme phenotypes. A. Presentation of the different layers used for multi-omic integration, the strategy leading to signature identification and schematic view of BD ranking. B. Heatmaps showing the mean of the Z-score distribution for each group, for all features selected in each layer. C. Plots showing individuals ranked by their predicted probability of belonging to the obese group, using models trained using data from individual layers, as well as a multi-layer predictive model (as indicated by the plot titles). Plots are ordered by decreasing log loss, with smaller values corresponding to better discrimination of individuals in the extreme phenotype group from all other individuals. D. Heatmap showing age and sex adjusted association values between (left) eigth prioritised lipid species and risk factors measured in the Fenland and present cohorts; and (right) a negative control set of five unselected lipid species and the same risk factors. Black asterisks indicate significant associations after correcting for multi-

DEG GO terms enrichment identified amongst the up-regulated pathways: ribosome formation, metabolism of amino acid and proteins, several immune related pathways and cytoplasm translation and amongst the down-regulated pathways: cholesterol metabolic process (through SREBF and miR33^59^) and mRNA processing pathways (**Table S18-S21**). We found genes whose expression was reduced in obese, to revert to the levels observed in the lean group: nine in macrophages (*RHPN1*, *DGKQ*, *TCTEX1D2*, *MVD*, *LDL-R*, *BCAR1*, *ANKRD33B*, *FASN*, *COL5A3*; overlap p value = 3.6*10^-8^, hyper-geometric test), seven in monocytes (*EPB41L3*, *LRRC8B*, *STARD4*, *ZNF331*, *SEMA6B*, *DSC2*, *RGPD8*; overlap p value = 5*10^-6^), five in neutrophils (*NAIP, RP11-1319K7.1, LINC01271, LINC01270, DNAH17*; overlap p value = 1.3*10^-5^) and ten in platelets (*CTC-429P9.4, XXbac-BPG300A18.13, RP11-386G11.10, MT-TG, TVP23C-CDRT4, SHE, MPZL3, CLIP1, RGPD1, RPL23AP7*; overlap p value=6.5*10^-5^). These indicate that lipoprotein metabolism (LDL-R), fatty acid synthesis (FASN) and cholesterol transport (STARD4) are restored after surgery. We also found two genes in macrophages (*SNHG5*, *EVI2A*; overlap p value = 0.03) and three in monocytes (*XXbac-BPG32J3.22, MEIS2, MS4A14*; overlap p value = 0.03) that move in the directions. While some genes, after bariatric surgery, reverted to expression level observed in lean individuals, the majority of DEG either did not revert to the values observed in lean individuals or were not differentially expressed in the comparison between the obese and lean groups. This suggests that the reduction in inflammatory signatures observed in these four cell types after bariatric surgery was achieved with the establishment, at least in the time frame investigated, of novel gene expression landscapes. Moreover, the overall small number of changes in DNA methylation observed, together with the short life span of the hematopoietic cells analysed, indicated that the change in exposure had little effect on the hematopoietic stem cell epigenome and that the effects observed in animal models^60^ were either species specific or were diluted and then lost with the turnover of the hematopoietic progenitor pool.

The effects of bariatric surgery at organism level were monitored with plasma proteomics. We quantified 3,098 plasma proteins; 604 of which were found to be differentially abundant (DAP; **Fig.3C** and **Table S24**) above ordinal Q-value of 1*10^-3^. Proteins whose levels increased after bariatric surgery (n=72) were enriched in GO terms related to tight junction and WNT, PI3K/AKT, sphingolipid signalling pathways. Proteins whose abundance decreased after surgery (n=532) were enriched in the following GO terms: cell cycle and DNA repair, ribosomal RNA metabolism and cell senescence, phagocytosis and T cell receptor signalling as well as FGF, IL2, VEGF and insulin signalling pathways (**Table S25**). Amongst these we also found NLRP3, a critical mediator of inflammation^61^ and several histones, normally released by cells undergoing apoptosis and netosis^62^. No changes in full blood count that could explain these changes were observed. We only noted an increase in mean platelet volume (p value = 0.03; paired t-test) and a reduction of the lymphocytes (p value = 0.03) and eosinophils (p value = 0.03; **Table S1**) counts. The plasma proteomic results showed that the changes after bariatric surgery were not limited to immune cells. To determine if any of them could be ascribed to a specific tissue, we determined which genes were tissue specific, using the GTEx project database^63^ (**Table S26;** METHODS). Tibia, coronary and aortic arteries, heart atrial appendage, heart left ventricle, and blood displayed an enrichment for tissue specific genes amongst DAP (p values: 1.6*10^-2^, 8*10^-3^, 2*10^-2^, 1.8*10^-2^, 1.6*10^-2^ and 5*10^-2^, respectively; hyper-geometric test; **Table S26**). Of the 13 blood specific genes encoding a DAP, six were also differentially expressed in at least one of the cell types (**Fig.3D**). These six genes have roles in immune response and leptin resistance^64^, immune pathways^65^, neutrophils recruitment during thrombosis^66^ and macrophage differentiation and inflammatory response^67^. The overall decrease observed indicated that vascular integrity, compromised by obesity^68^, was restored, as also observed by Albrechtsen and colleagues^69^.

Furthermore, monocytes and macrophages data allowed us to explore the effect of bariatric surgery on trained immunity^53^, which has been shown to play a role in atherosclerosis^70, 71^. Genes displaying an active promoter (H3K4me3), with or without β-glucan treatment, significantly overlapped with DEG in the obese versus post surgery comparison (p value = 4.5*10^-2^ and p value = 7.7*10^-3^**; Table S27**). This suggested that bariatric surgery had a positive impact on innate immune cells, and indicated that trained immunity acts downstream of the hematopoietic stem cell pool and its effects were diluted and eventually lost with the renewal of the hematopoietic progenitors pool.

To determine the impact of the changes observed at molecular levels on the functional phenotypes of these cell types, we performed functional tests on neutrophils and platelets. After bariatric surgery, neutrophils showed a reduction in their ability to adhere both when unstimulated (**Fig. 3E**), as well as, when subjected to a variety of stimuli (DTT, LBP, PAM3, PAF and fMLP; **Fig. 3F**), but not when treated with TNFalpha or PMA. These results were accompanied by a reduction in the cell surface levels of CD16 and CD32, but not CD66b, CD63, CD62L or CD11b (paired t-test, all result in **Table S28**). Alongside, we also performed platelet functional tests, which showed a reduction in P-selectin surface exposure upon collagen stimulation, but not upon ADP or thrombin stimulation. These results were accompanied by a reduction in the cell surface levels of fibrinogen receptor (CD61 and CD41b) and CD36, the thrombospondin receptor that acts as scavenger for oxidized LDL. No changes were observed for CD49b, CD42a, CD42b, CD29 and CD9 (paired t-test, all results in **Table S28**).

The integration of gene expression and proteomic data showed that some of the changes at transcriptional level were directly involved in the reduction of the proinflammatory environment, but also highlighted a conspicuous involvement of other levels of regulation. Notably, we collected across several data layers evidence to suggest a diminished ability of the cells to use neutrophil extracellular traps (NETosis)^72^ after bariatric surgery (**Fig. 4**). NETs are formed by chromatin (DNA and histones), granular antimicrobial proteins and cytoplasmic proteins, are normally found at low levels in the circulation^73^, however in the presence of pathogens or sterile inflammation, such as the increase of reactive oxygen species observed in obese individuals^74^, NETs levels are increased. We observed decrease plasma levels of NLRP3 a critical mediator of the inflammasome^61^, RAC2 a protein directly involved in NETs promotion^75^, MYO1G, a protein promoting immune cells interaction^76^ and several histones, core component of the chromatin released during NETosis^76^ (all in **Fig. 3C**). Additionally, the upregulated genes in obese individuals indicated increased activity of neutrophils and monocytes (**Table S19-S20**).

**Figure 4.**
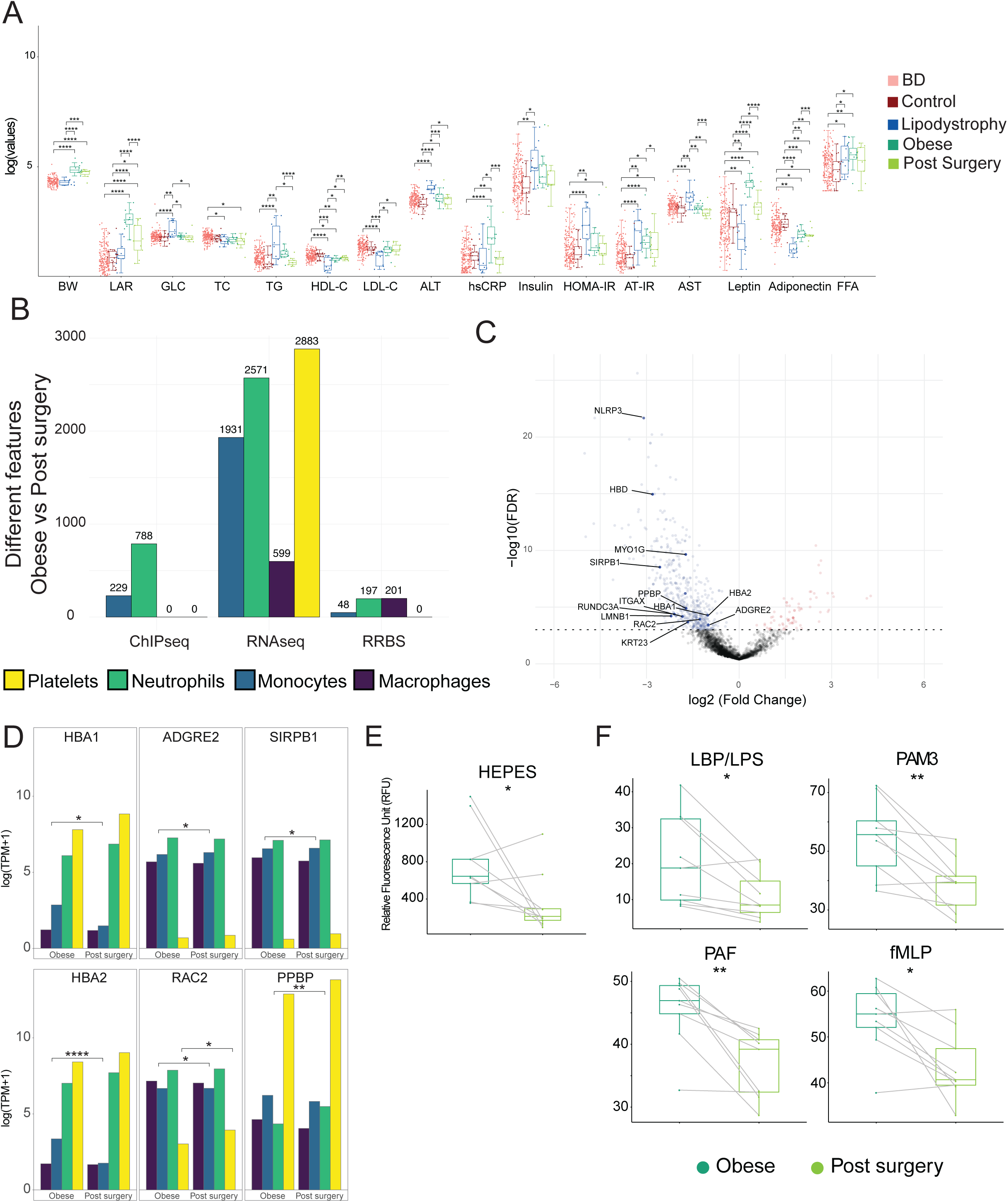
Effect of bariatric surgery on transcriptional profile, epigenetic landscape and cell functions. A. Body weight (BW) and biochemical values distribution across the four studied groups: obese (dark green); lipodystrophy (blue); blood donors (BD) (light red); and post bariatric surgery patients (light green). Asterisks indicate result of significance from multiple logistic regression models and conditional multiple logistic regression for obese versus post surgery comparison. Significance is annotated as follows: * P≤ 0.05, ** P ≤ 0.01, *** P ≤ 0.001, **** P ≤ 0.0001. B. Bar Plot shows number of features significantly different when comparing obese individuals before and after bariatric surgery, colored by cell types. C. Volcano plot showing differentially abundant plasma proteins when comparing obese individuals before and after bariatric surgery. Whole blood specific genes associated with differentially abundant proteins have been annotated. D. RNA-Seq expression in the 4 different cell types of highlighted proteins in C. Asterisks indicate if the gene was differentially expressed in at least one cell type. E. Neutrophil ability to attach in the absence of any stimuli after bariatric surgery and it is expressed using the plate reader arbitrary units (RFU) . F. Adhesion percentage of neutrophils measured in the presence of different pro-inflammatory molecules in obese (dark green) and post surgery (light green) individuals. Asterisks indicate the result of significance from paired t-test. Significance is annotated as follows: * P≤ 0.05, ** P ≤ 0.01.

**Figure 5.**
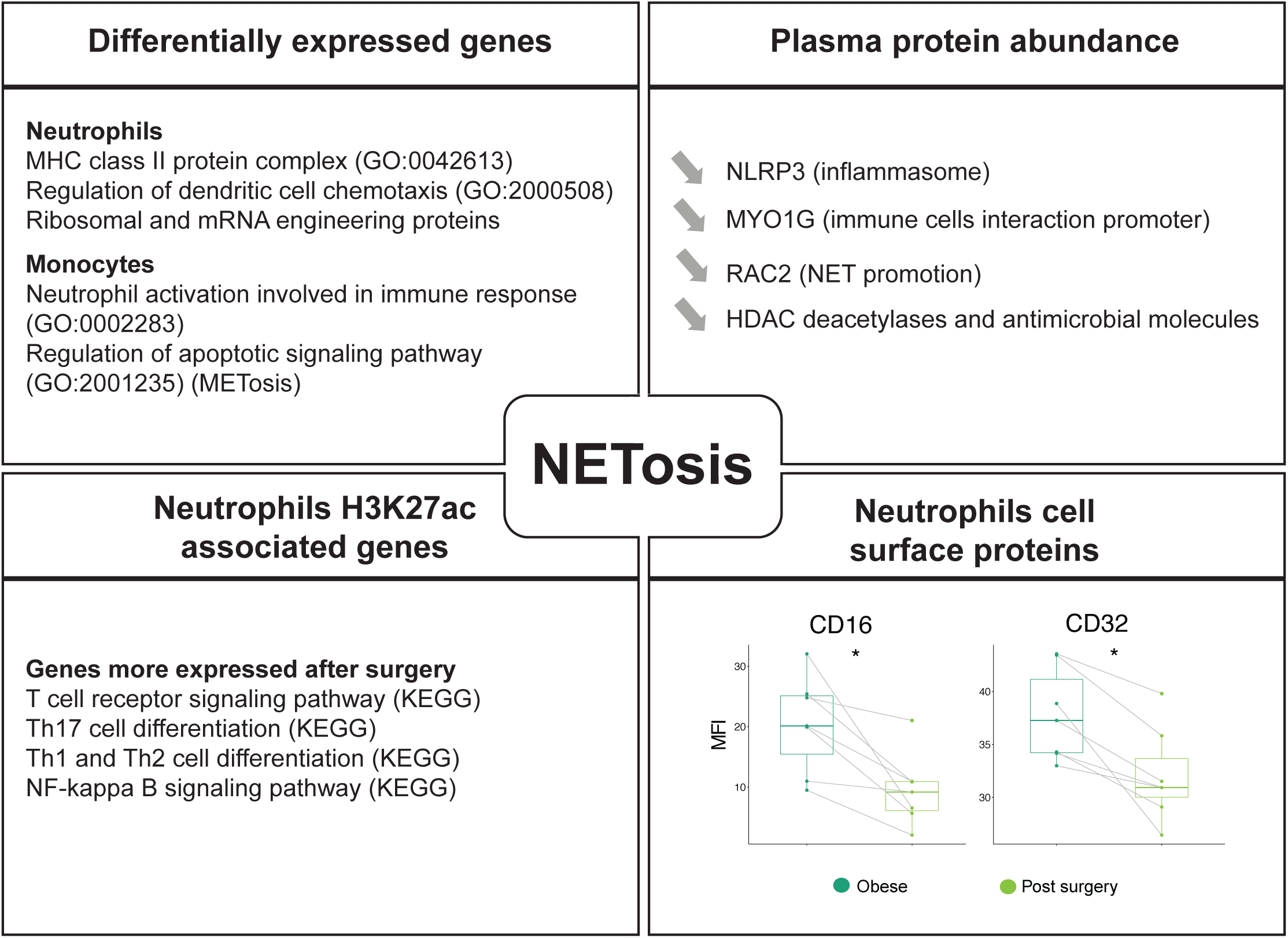
Different omic layers contribution to NETosis reduction 6 months after bariatric surgery in mordibly obese individuals. Significance is annotated as follows: * P≤ 0.05, ** P ≤ 0.01, *** P ≤ 0.001, **** P ≤ 0.0001.

Lastly, we also observed a decrease in the ability of neutrophils to adhere, alongside changes in their surface protein levels, such as CD16 and CDE32, also previously associated with NETosis^77^ (**Table S28**). In addition, genes associated with DAcR in neutrophils post surgery showed enrichment in the T-cell receptor signaling pathway, in particular Th17 cell differentiation (**Table S22**), which suggested a restored ability for neutrophils to activate T-cell through NETosis.

The results of the comparisons between lipodystrophy and post bariatric surgery and post bariatric surgery and lean (**Fig.S2**) are available in **Tables S8 to S17**.

Altogether, these findings indicate that many key players in thrombus formation have reduced ability to respond to stimuli after bariatric surgery. Our analyses showed that the extreme phenotype groups could be separated using each of the different layers of information. Given the limited number of changes identified using single layer univariate comparisons, we sought to identify multivariable signatures by combining information across layers to better characterise and discriminate between the extreme phenotype and lean groups.

### Multi-omic signature classification of extreme phenotypes

Six lean (METHODS) and six obese individuals, for which we had complete measurements on all layers, in monocytes and in neutrophils, were used to define training sets. Since multivariable selection approaches have provided an effective means to integrate multiple omics layers and elucidate disease signatures^77, 78^, we applied elastic net penalised logistic regression^80^ to identify signatures associated with an increased probability of belonging to the obese group and therefore to have some or all features associated with CMS (**Fig.5A**). We performed this analysis independently for each data layer (METHODS). The variables selected into each signature defined patterns characterising the groups (**Fig.5B**; **Table S29-31**) and the biometric variables were then used to construct multivariable logistic regression models, using either the variables selected from a single data layer or all selected variables, across omic layers (METHODS). All models, single layer or multi-layer trained, allowed us to rank individuals according to their probability of belonging to the obese group (**Fig.5C** and **Fig.S3A**). We quantified the log loss^82^ (or cross-entropy loss; **Fig.5C** and **Fig.S3A**) and demonstrated that the multi-layer model provided the greatest separation, followed by the models trained on the RNA-seq, then those trained on metabolites and methylation. Ranking all individuals by their probability of belonging to the obese group according to the multi-layer model showed that the obese and lipodystrophy individuals had the highest probabilities of belonging to the obese group, with 8/10 lipodystrophy individuals predicted to belong to the obese group with probability >0.5. Qualitatively similar results were obtained when using the lean and lipodystrophy individuals as training data (**Fig S3B**), suggesting that individuals belonging to the extreme phenotype groups tend to be more similar to one another, when taking into account all of the data layers, than to individuals outside these groups. We moreover note that amongst the 20 BD predicted by the multi-layer model to belong to the obese group with highest probability, 4 were in the lowest quartile for weight and 8 were in the lowest quartile for LDL (**Table S32**), demonstrating that our multi-omic model is not simply recapitulating features of classic CMS presentation. The differences observed when comparing the models trained using the lipodystrophy instead of the obese group likely reflect the higher heterogeneity in the latter, lacking a high penetrance genetic cause. External cohorts with similar data layers will be required to establish the predictive utility of our models, and to fully validate the omic signatures identified in this study. However, because lipidomic data from external cohorts were available, we focused on the validation of the lipidomic signature. We prioritised a reduced set of nine lipid species from the signature to test for univariate association with known CMS risk factors (METHODS, **Fig.5D**), including eight matched lipid species measured in a subset of 1,507 participants in the Fenland study^83^. After correcting for multiple testing, 61% (225/368) of associations remained significant. Triacylglycerol 52:2 and 50:1 were positively associated with several risks factors (fasting plasma glucose, fasting insulin level, HOMA-IR, a fatty liver index, HbA1c, leptin, LDL-C, hsCRP, TG, BMI, fat mass, ALT, and ferritin; **Table S33**) and inversely associated with adiponectin and HDL-C. Phosphatidylcholine (40:7), (38:7), (38:6), (35:2) and O (36:2) were inversely associated with all factors except for adiponectin and HDL-Further supporting our findings, phosphatidylcholine (38:6 and 36:2) had previously been identified in obesity studies^84^; and, triacylglycerol (50:1 and 52:2) had previously been linked to NAFLD^83^ and NASH^85^. To assess the specificity of the results, we repeated the analysis with five lipid species randomly selected (METHODS) from those not included in the signature. Only 21% of associations were significant (49 out of 230 tests). The same pattern of associations was also found in our study (**Fig.5D; Table S33**), as well as, in a biopsy-confirmed non-alcoholic steatohepatitis (NASH) cohort comprising 73 individuals^85^ (**Fig.S4**; **Table S33**). We showed the diagnostic value of the prioritised lipid species through their association with major cardiometabolic risk factors in the Fenland study and in the present study; as well as, albeit not significantly due to the small sample size, in the NASH cohort.

## Conclusion

Our overall goal was to develop an integrative multi-omic strategy to combine information collected across different -omics layers in order to account for the impact of genetic and environmental differences on each of them. We generated data from extreme metabolic phenotype groups to obtain a signature for CMS and then used this signature to determine the cardiometabolic status of a group of individuals (BD) that due to age, are at increased risk of developing CMS. Substantial annotations in our analysis identified the reduction of inflammation and the reduction of the ability to form extracellular traps as key consequences of bariatric surgery in innate immune cells and platelets. Further investigations of the molecular basis underlying the priming of these innate immune cells will help to understand which features, such as small molecules or metabolites, promote abnormal inflammation and extracellular traps formation, providing possible avenues for future clinical treatments.

## Supporting information

Extended Methods Metabolon

## Acknowledgments and funding

L.S. is supported as PhD student by British Heart Foundation Cambridge Centre of Excellence; M.C.S is supported by a MRC Clinical Research Training Fellowships (MR/R002363/1); D.B.S is supported by the Wellcome Trust (WT 107064), the MRC Metabolic Disease Unit (MRC_MC_UU_12012.1), and The National Institute for Health Research (NIHR) Cambridge Biomedical Research Centre and NIHR Rare Disease Translational Research Collaboration; K.D. is supported as a HSST trainee by NHS Health Education England; P.D.W.K is supported by the Medical Research Council (MC_UU_00002/13). The Human Research Tissue Bank is supported by the NIHR Cambridge Biomedical Research Centre. M.F. is supported by the British Heart Foundation (FS/18/53/33863). D.S. work has been supported in part by an Isaac Newton fellowship to M.F, L.L.N. is supported by the NIHR Leicester Biomedical Research Centre and the John and Lucille Van Geest Foundation.

## Author contributions

Conceptualization: D.S., J.J.L, L.G., L.L.N., M.V., P.D.W.K. and M.F.; Methodology: D.S., A.C., T.H.C., L.L.N., P.D.W.K. and M.F.; Patients recruitment, samples collection & data generation: J.J.L., F.B., S.F, K.R., A.F., H.M., J.B., C.K., C.L.A., A.K., L.L.N., O.S., P.A.Q., L.S., M.C.S., D.B.S., C.L., C.B., K.D., G.M., M.A., M.V., A.P., D.B.S. and M.F.; Software and algorithm implementation: D.S., A.C., B.E. and P.D.W.K.; Data curation: D.S., A.C., C.L.A., L.L.N. and P.D.W.K; Data analysis, D.S., A.C., T.H.C., B.E., L.L.N., K.D. and P.D.W.K; Investigation, D.S., A.C., T.H.C., P.A.Q., L.L.N., A.P., S. d’A., M.V., P.D.W.K. and M.F.; Fenland cohort management and analysis: N.J.W., M.P and C.L.; Writing groups – Original Draft: D.S., A.C., T.H.C., L.L.N., P.D.W.K. and M.F.; Writing – Review & Editing, everyone; Figures preparation: D.S., A.C., M.P, P.D.W.K. and M.F.; Funding Acquisition, M.F.; Supervision, J.L.G., C.B., N.J.W., C.L., L.L.N. P.D.W.K. and M.F., Project administration, M.F.

## Methods

### Data availability

The datasets generated during this study are available at EGA under study ID EGAS00001003780.

The codes generated during this study and all supplementary tables are available at GitLab https://gitlab.com/dseyres/extremephenotype.

### Patients recruitment and ethics

Obese individuals referred for obese surgery by the obesity clinic and lipodystrophy patient cared for by the National Severe Insulin Resistance Service respectively, both based at Addenbrooke’s hospital, Cambridge University Hospitals were recruited to this study together with healthy individuals. Informed consent was obtained under the “Inherited Platelet Disorders” ethics (REC approval 10/H0304/66 for patients and 10/H0304/65 for healthy controls, NRES Committee East of England-Cambridge East).

BluePrint work package 10 (WP10) volunteers (representing the blood donors, “BD”, cohort) were recruited amongst NHS Blood and Transplant donors after informed consent under the “A Blueprint of Blood Cells” ethical approval (REC approval 12/EE/0040 NRES Committee East of England-Hertfordshire).

“BioNASH” Cohort consisted of 73 consecutive patients recruited at the NASH Service at the Cambridge University Hospital. All the patients had a clinical diagnosis of NAFLD (patients with alternate diagnoses and etiologies were excluded) and histology scored. This study was approved by the local Ethics Committee; all patients gave their informed consent for the use of data (Biochemistry and clinical history) and samples for research purposes. The principles of the Declaration of Helsinki were followed.

### Cell types isolation

Whole blood (50ml) in citrate tubes was obtained after informed consent. Platelet rich plasma (PRP) was separated from the cellular fraction by centrifugation (20’, 150g and very gentle break) for platelet isolation. Platelets were then isolated from PRP after 2 more spins as above and leukodepleted using anti CD45 Dynabeads (Thermofisher) following the manufacturer’s instructions. Purified platelets were stored in TRIzol (Invitrogen) until RNA extraction. The remaining cells were resuspended in buffer 1 and separated on a Percoll gradient. Neutrophils were harvested from the red blood cell pellet after red cell lysis (4.15 g NH4Cl, 0.5 g KHCO3 and 18.5 mg EDTA (triplex III, 0.01%) in 500 ml of water) and aliquots prepared for RNA extraction (TRIzol), DNA extraction for RRBS (snap frozen pellet) and ChIP-Seq (formaldehyde fixation, see below). Monocytes were isolated from the peripheral blood mononuclear cell (PBMC) layer by CD14 positive selection (Miltenyi) and aliquots prepared for RNA extraction (TRIzol), DNA extraction for RRBS (snap frozen pellet) and ChIP-Seq (formaldehyde fixation, see below). Macrophages were cultured by plating 14*106 PBMC resuspended in 2 ml macrophage media (Macrophage-SFM [with L-Glutamine without Antibiotics], Fisher Scientific UK LTD). After 1h 30’ non adherent cells were removed and 1 ml fresh macrophage media added together with 400 *μ*l of autologous serum. Culture media was replaced after 3 or 4 days. On day 7 cells were harvested for RNA extraction (TRIzol), DNA extraction for RRBS (snap frozen pellet) and ChIP-Seq (formaldehyde fixation). Cell purity was determined by flow cytometry as follows: neutrophils CD66b (BIRMA17c, FITC, 9453 https://ibgrl.blood.co.uk/), CD16 (VEP13, PE, 130-091-245 Miltenyi) and CD45 (HI30, PE-CY5.5, MHCD4518 Invitrogen); monocytes CD14 (MφP9, FITC, 345784 BD), CD16 (B73.1 / leu11c, PE, 332779 BD), CD64(10.1, PerCP-Cy5.5, 561194 BD), CD45 (HI30, PE-CY7, MHCD4512 Invitrogen); macrophages panel 1: CCR7/CD197 (150503, FITC 561271 BD), CD25-PE MACS 120-001-311 (10ul/test), CD14 (TuK4, PE-Cy5.5, MHCD1418 Invitrogen), CD40 (5C3, PE-Cy7, 561215 BD). Panel 2: CD206 (19.2, PE, 555954 BD), CD36 (SM***Φ***, FITC, 9605-02 Southern Biotech), CD45 (HI30, PE-Cy5.5, MHCD4518 Invitrogen). Samples whose purity was below 90% were discarded. BD samples isolation has been extensively described in Chen et al..

### RNA sequencing

#### RNA extraction

RNA extraction from samples stored in TRIzol was performed following the manufacturer’s instructions. Briefly, tubes were retrieved in small batches and thawed on ice. Prior to extraction samples were vortexed for 30” to ensure complete lysis and let for 5’ at room temperature. Samples were then transferred to heavy phase lock tubes (5prime) to separate RNA in the aqueous phase from the organic phase. RNA was precipitated from the former with isopropanol and glycogen. The RNA pellet was resuspended in RNase free water. Purified RNA was stored in single use aliquots. Each sample was quality controlled by a Bioanalayser (Agilent) and quantified via Qubit (Thermofisher).

#### Library preparation and sequencing

For cell types isolated from obese and lipodystrophy patients and day controls we used 100 ng of total RNA for neutrophils, monocytes and macrophages and 200 ng for platelets. o libraries were prepared for sequencing using the Kapa stranded RNA-Seq kit with riboerase (Roche) according to the manufacturer’s instructions and sequenced 150bp paired end on Illumina HiSeq 2500 or Illumina HiSeq 4000. BD RNA-Seq data (extensively described in Chen et al.^43^) were retrieved from European Genome-phenome Archive (EGA) - EMBL-EBI after application to the Data Access Committee.

#### Quantification

FastQ files were first checked for sequencing quality using FastQC (v.0.11.2) [https://www.bioinformatics.babraham.ac.uk/projects/fastqc/] and quality trimmed with TrimGalore! (v.0.3.7) [https://www.bioinformatics.babraham.ac.uk/projects/trim_galore/]. Transcript-level abundance was estimated using Kallisto (v0.42)^86^ with 100 bootstrap iterations in single-end mode for extreme phenotype samples in order to minimize technical batch effect with BD cohort. Transcript abundances were then summarized to gene-level with Tximport R package (v1.9) ^87^ by using tximport function and Ensembl reference transcriptome (Ensembl Genes 96)^87, 88^. This step provides an input count matrix for DESeq2 (v.1.21.21)^89^. DESeq2 was used to normalize counts by library size and transformed by variance stabilisation (VST). We corrected for sequencing batch effects by using Combat (from sva R package (v.3.29.1))^89, 90^ and individual status as covariate. Non-autosomal genes and those with no or low variance across individuals were removed. The final gene sets (including coding and non-coding genes) were formed of 10,925 genes for monocytes and of 26,634 for neutrophils. Quality metrics are reported in **table S4**.

#### Differential analysis

For differential analysis, transcript-level abundance was estimated by Kallisto with 100 bootstrap iterations in paired-end mode for each group (obese, post surgery, lipodystrophy patients and lean individuals) using Ensembl reference transcriptome (Ensembl Genes 96). Transcript abundances were then summarized to gene-level with Tximport R package (v1.9) by using tximport function and DESeq2 object was created using DESeqDataSetFromTximport function from DESeq2 R package (v.1.21.21). Differential analysis was performed using the DESeq function from DESeq2 and we used age and gender as covariates. Log fold changes were corrected with the lfcShrink function from DESeq2. Genes with FDR < 5% were marked as differentially expressed. Lean individuals were selected from control group by applying the following criteria: BMI < 25, glycaemia (GLUC) <5.4 mmol/L, TG <1.7 mmol/L, LDL <2.59 mmol/L, HDL >1 mmol/L for men and >1.3 mmol/L for women, HOMA score< 2.2. For obese versus post surgery comparison, we considered only paired samples ([S01RS6;S022QS][S01Y9G;S022UK][S01WCI;S0232Z][S01TEQ;S0234V][S01WXD;S02 3EB][S01WFC;S023F9][S01Y7K;S023H5][S022TM;S023PQ][S01XJ0;S023RM][S01SYR; S0240Z][S022GB;S0245P]) and therefore performed a paired analysis by adding relationship information as covariate in the design formula.

For each cell-type, functional annotation was performed with genes differentially expressed in each comparisons, taking into account fold change direction. Lists of genes were submitted to EnrichR using the R package EnrichR (v.1.0) ^91, 92^ and the following databases: BioCarta_2016, DSigDB, GO_Biological_Process_2018, GO_Cellular_Component_2018, GO_Molecular_Function_2018, HMDB_Metabolites, KEGG_2019_Human, Reactome_2016 and WikiPathways_2015.

### Chromatin Immunoprecipitation sequencing

#### Sample preparation

Cells were fixed immediately after purification with 1% w/v formaldehyde for 10 min and quenched using 125 mM glycine before washing with PBS. Samples were sonicated using a Bioruptor (Diagenode), final SDS concentration of 0.1% w/v for 9 cycles of 30 s ‘on’ and 30 s ‘off’, and immunoprecipitated using an IP-Star Compact Automated System (Diagenode) using the histone H3K27ac antibody C15410196 (lot 1723-0041D) Diagenode. Immunoprecipitated and input DNA were reverse cross-linked (65 C for 4 h), treated with RNase and Proteinase K (65 C for 30 min).

#### Library preparation and sequencing

DNA was recovered with Concentrator 5 columns (Zymo) and prepared for sequencing using MicroPlex Library Preparation Kit v2 (C05010012, Diagenode). Libraries analysed using High Sensitivity Bioanalyzer chips (5,067–4,626, Agilent), quantified using qPCR Library Quantification Kit (KK4824, Kapa Biosystems), pooled and sequenced with a 50bp single end protocol on Illumina HiSeq 2500 or Illumina HiSeq 4000.

#### Peak calling and quantification

FastQ files were first checked for sequencing quality using FastQC (v.0.11.2) and quality trimming was applied on reads with TrimGalore! (v.0.3.7). Trimmed FASTQ files were aligned to the human genome (Ensembl GRCh38.80) with BWA (v.0.7.12)^93^ *aln* and *samse* functions with default parameters. Low mapping quality reads (-q 15), multi-mapped and duplicate reads were marked and removed with respectively samtools (v.1.3.1)^94^ and picard (http://broadinstitute.github.io/picard v.2.0.1).

A combination of quality metrics was used to assess sample quality: number of uniquely mapped reads, number of called peaks, NSC (Normalized strand cross-correlation) and RSC (relative strand cross-correlation) computed with Phantompeakqualtools (v.1.2)^95, 96^, area under the curve (AUC), X-intercept and Elbow Point computed with plotFingerPrint function from deepTools suite (v.3.0.2)^97^ with --skipZeros --numberOfSamples 50000 options. Peaks were called with MACS2 (v.2.1.1) with --nomodel --shift -100 --extsize 200, a qvalue threshold of 1e-3 options and celltype matching input file scaled to sample read number. We used the MACS2 randsample function to downscale inputs. We then computed a score by summing values obtained for each range of these metrics. We applied a threshold of -3 (total) to select the best quality data.

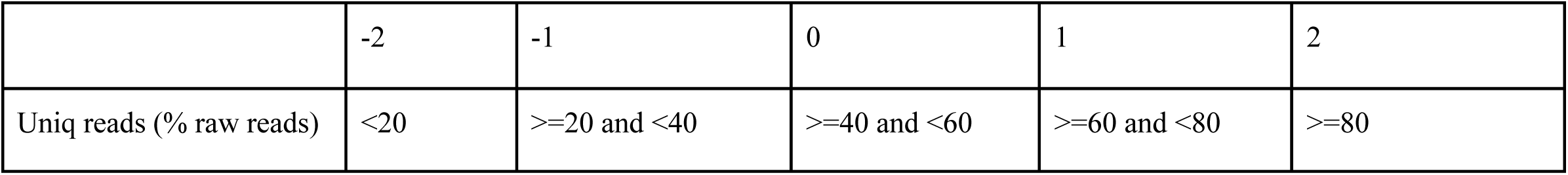
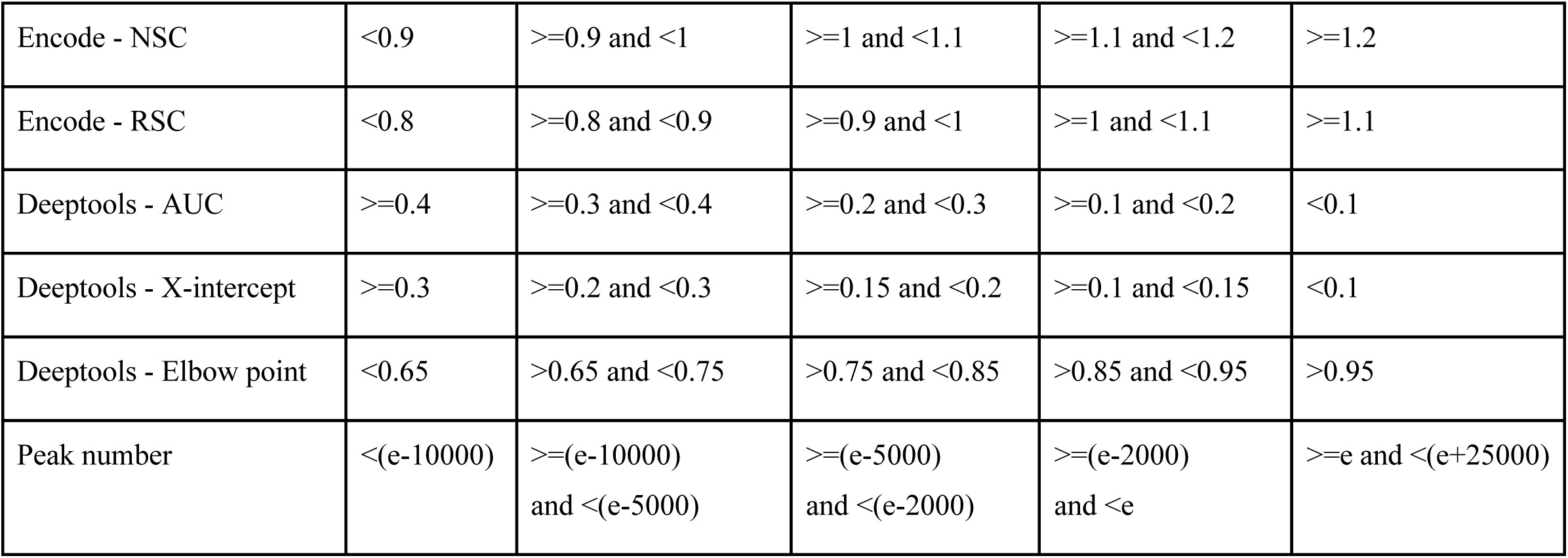

To build the ChIP-Seq layer for integrative analysis, we defined a master set of peaks and quantified H3K27ac ChIP-Seq signals under these peaks. Peaks shared by at least 5 individuals were merged using R package DiffBind (v2.9)^98^. We obtained 67,763 and 49,188 peaks for monocytes and neutrophils, respectively. Minimum merged peak size was 244bp and 235bp, median peak size 1,392bp and 1,648bp and maximum peak size 75,534bp and 60,528bp for monocytes and neutrophils, respectively. We didn’t filter out very large merged peaks as they represent less than 3% of total peaks and indicate large acetylated regions. Read counts under merged peaks were TMM normalized using effective library size and logit transformed into count per million (CPM). Sequencing center batch effect was corrected with Combat (from sva R package (v.3.29.1)) using individual status (Patient/Donor) as covariate. Non-autosomal and no or low variance peaks across individuals were removed. The final master set of peaks counted 25,595 regions in monocytes and 26,300 regions in neutrophils. Quality metrics are reported in **table S3**.

#### Differential analysis

For differential analysis, we used DiffBind with the built-in DESeq2 method for statistical analysis. We merged peaks present in at least 50% of individuals and asked that all individuals have a FRiP value (Fraction of Reads in Peaks) over 5%. We then applied a FDR threshold of 5% to select H3K27ac peaks differentially acetylated peaks. We used age and gender as covariates. For obese versus post surgery comparison, we considered only paired samples and therefore performed a paired analysis by using the block factor in DESeq2. Differentially acetylated regions (DAcR) were annotated with HOMER (v.4.10)^99^, annotatePeak function and Hg38 RefSeq genome annotation (http://homer.ucsd.edu/homer/data/genomes/hg38.v6.0.zip).

Functional annotation was performed on genes within a window of 10kb around each DAcR, taking into account fold change direction. Similarly to RNA-Seq, lists of genes were submitted to EnrichR interrogating the same databases. Annotation results are available in **table S22**.

### Illumina 450K arrays and reduced representation bisulfite sequencing (RRBS)

#### Arrays and libraries preparation and sequencing

BD Infinium Human Methylation 450 arrays (Illumina) were retrieved from the European Genome-phenome Archive (EGA) - EMBL-EBI. DNA extraction and array generation have been described in detail in Chen et al.^43^. Briefly, cells were lysed using guanidine hydrochloride, sodium acetate and protease lysis buffer. DNA was extracted using chloroform and precipitated in ethanol prior to washing and resuspension in ultra-pure water. 500ng of DNA for each monocyte and neutrophil sample was randomly dispensed onto a 96-well plate to reduce batch effects. Samples were bisulfite-converted using an EZ-96 DNA Methylation MagPrep Kit (Zymo Research) following the manufacturer’s instructions with optimized incubation conditions (i.e., 16 cycles of 95C for 30 s, 50C for 60 min; followed by 4C until further processing). Purified bisulfite-treated DNA was eluted in 15 mL of M-Elution Buffer (Zymo Research). DNA methylation levels were measured using Infinium Human Methylation 450 arrays (Illumina) according to the manufacturer’s protocol.

For RRBS, 100 ng of genomic DNA were digested for 6h at 65°C with 20 U TaqI (New England Biolabs) and 6h hours at 37°C with 20 U of MspI (New England Biolabs) in 30 μl of 1x NEBuffer 2. To retain even the smallest fragments and to minimize the loss of material, end preparation and adaptor ligation were performed in a single-tube setup. End fill-in and A-tailing were performed by addition of Klenow Fragment 3’ --> 5’ exo-(Enzymatics) and dNTP mix (10 mM dATP, 1 mM dCTP, 1 mM dGTP New England Biolabs). After ligation to methylated Illumina TruSeq LT v2 adaptors using T4 DNA Ligase rapid (Enzymatics), the libraries were size selected by performing a 0.75x clean-up with AMPure XP beads (Beckman Coulter). The libraries were pooled based on qPCR data and subjected to bisulfite conversion using the EZ DNA Methylation Direct Kit (Zymo Research) with changes to the manufacturer’s protocol: conversion reagent was used at 0.9x concentration, incubation performed for 20 cycles of 1 min at 95°C, 10 min at 60°C and the desulphonation time was extended to 30 min. These changes increase the number of CpG dinucleotides covered, by reducing double-strand break formation in larger library fragments. Bisulfite-converted libraries were enriched KAPA HiFi HS Uracil+ RM (Roche). The minimum number of enrichment cycles was estimated based on a qPCR experiment. After a 1x AMPure XP clean-up, library concentrations were quantified with the Qubit Fluorometric Quantitation system (Life Technologies) and the size distribution was assessed using the Bioanalyzer High Sensitivity DNA Kit (Agilent).

#### Processing and quantification

All Infinium Human Methylation 450 array data pre-processing steps were carried out using established analytical methods incorporated in the R package RnBeads (v.1.13.4)^100^. First, we performed background correction and dye-bias normalization using NOOB^101^, followed by normalization between Infinium probe types with SWAN^102^. Next, we filtered out probes based on the following criteria: median detection p value 0.01 in one or more samples; bead count of less than three in at least 5% of samples; ambiguous genomic locations^103^; cross-reactive and SNP-overlapping probes^104^.

The RRBS samples were sequenced on Illumina HiSeq3000 platform in 50bp single-end mode. Base calling was performed by Illumina Real Time Analysis (v2.7.7) software and the base calls were converted to short reads using Illumina2bam (1.17.3 https://github.com/wtsi-npg/illumina2bam) tool before de-multiplexing (BamIndexDecoder) into individual, sample-specific BAM files. Trimmomatic (v0.32)^105^ was used for trimming the adapter sequences. Trimmed short read sequences were aligned onto the GRCh38/hg38 human reference genome with BSMAP(v2.90)^106^ aligner in RRBS mode which was optimized for aligning the RRBS data while being aware of the restriction sites and with the following options: -D C-CGG -D T-CGA -w 100 -v 0.08 -r 1 -p 4 -n 0 -s 12 -S 0 -f 5 -q 0 -u -V 2. R package RnBeads was used to filter out low confidence sites: sites overlapping any SNP, having a coverage lower than 5 and high coverage or missing in more than 5% or individuals were filtered out. Integration analysis required to attenuate technology effect between 450K arrays and RRBS. To this goal, we generated RRBS data for 14 BluePrint donors for which we already have 450K array data in monocytes, and 9 in neutrophils. We first removed non reproducible sites between technologies as follows: for monocytes and neutrophils, 1) liftover 450K sites to Hg38 using UCSC liftover tool^107^, 2) keep overlapping sites between array and RRBS, 3) filter out sites with high variation in methylation percentage observed in more than 70% of individuals. We excluded 844 and 1,127 sites for monocytes and neutrophils respectively. We have also excluded sites on sex chromosomes and imputed missing values using KNN networks (impute.knn function from impute R package (v.1.55.0)) [Hastie T, Tibshirani R, Narasimhan B, Chu G (2019). impute: impute: Imputation for microarray data.] with 10 nearest neighbors. Finally, we adjusted for batch effects using an empirical Bayesian framework, as implemented in the ComBat function of the R package SVA (v.3.29.1) and individual status as covariate, transformed beta values to M values using beta2m function in R package lumi (v.2.33.0)^108, 109^, normalize by quantile using normalize.quantiles function from R package preprocessCore (v.1.43.0) [Bolstad B (2019). preprocessCore: A collection of pre-processing functions.] and remove zero or low variance sites. The final data matrix used for multi-omic integration, comprised DNA methylation M-values across 24,311 CpG sites and 210 samples in monocytes and 24,217 CpG sites and 203 samples in neutrophils.Quality metrics are reported in **table S5** and **S6**.

#### Differential analysis

For differential analysis, we used the methylKit R package (v.1.8.1)^110^ and we compared only RRBS data. We first extracted methylation ratios from BSMAP mapping results with methratio.py python script provided with BSMAP. We then removed all sex chromosomes sites and filtered out non-retained sites from RnBeads RRBS processing. Finally, we used the methRead function from methylKit R package in CpGs context at base resolution to read in the input files and calculateDiffMeth function correcting for overdispersion (overdispersion="MN") and applying Chisq-test. We used age and gender as covariates. Q Values are then computed using the SLIM method^110, 111^. We applied two thresholds: difference of methylation > 25 and qvalue < 0.05 and retrieved differentially methylated sites (DMS) with getMethylDiff function specifying type=”hypo” or type=”hyper” option to get down and up methylated CpGs respectively.

For obese (pre) versus post surgery comparison, we considered only paired samples and therefore performed a paired analysis. DMS were annotated with HOMER (v.4.10), annotatePeak function and Hg38 RefSeq genome annotation (http://homer.ucsd.edu/homer/data/genomes/hg38.v6.0.zip).

Functional annotation was performed on genes within a window of 10kb around each DMS, taking into account fold change direction. Similarly to RNA-Seq and ChIP-Seq, lists of genes were submitted to EnrichR interrogating the same databases. Annotation results are available in **table S23**.

### Plasma biochemistry assays

Plasma biochemistry assays were performed in the Core Biochemical Assay Laboratory, Cambridge University Hospitals (https://www.cuh.nhs.uk/core-biochemical-assay-laboratory) as described in supplementary material and methods. Homeostatic Model Assessment for Insulin Resistance (HOMA) score as follows: (glucose (mg/dL) x insulin (mIU/L)) / 405, and adipose tissue insulin resistance (AT) score as follows: insulin (µU/mL) x free fatty acids (mmol/L).

### Plasma metabolites measurement

#### Metabolites quantification

Metabolites profiling of obese and lipodhystrophy patients, day controls and blood donors (BD participants) was performed by Metabolon Inc. (https://www.metabolon.com/) using their standard protocol. Briefly, Metabolon analytical platform incorporates two separate ultra-high performance liquid chromatography/tandem mass spectrometry (UHPLC/MS/MS2) injections and one gas chromatography GC/MS injection per sample. The UHPLC injections are optimized for basic species and acidic species. The numbers of compounds of known structural identity (named biochemicals) as well as compounds of unknown structural identity (unnamed biochemicals) detected by this integrated platform were respectively of 793 and 362 for the first batch and 947 and 433 for the second batch (with an overlap of 786 and 359 compounds respectively). All samples were rescaled to set the median to 1, missing values were imputed using KNN networks (impute.knn function from impute R package (v.1.55.0) with the following options: number of nearest neighbors=10, maximum missing values per metabolites < 50% and maximum missing values for individuals < 80%.) Finally, we adjusted for batch effects using the ComBat function of the R package SVA (v.3.29.1) and individual status as covariate.

### Plasma lipids measurement

Plasma was frozen in dry ice immediately after collection and stored at -80C until analysis. Samples were prepared essentially as previously described^112^. Briefly, a 15 μL sample, controls and blanks were placed in a predefined random order across 96-well plates (Plate+, Esslab, Hadleigh, UK). To which, 750 µL methyl tert-butyl ether was added, along with 150 µl of internal standard mix, containing the following six internal standards (IS): 1,2-di-o-octadecyl-sn-glycero-3-phosphocholine (0.6 µM), 1,2-di-O-phytanyl-sn-glycero-3-phosphoethanolamine (1.2 µM), C8-ceramide (0.6 µM), N-heptadecanoyl-D-erythro-sphingosylphosphorylcholine (0.6µM), undecanoic acid (0.6µM), and trilaurin (0.6 µM), (Avanti Polar Lipids and SIgma Aldrich). Quality controls were derived from pooling all samples and serially diluting with chloroform. 25 µl of the sample/IS mixture was transferred to a glass coated 384 well plate and 90µl mass spectrometry (MS) mix [7.5mM NH4Ac IPA:MeOH (2:1)] added and then sealed. Lipidomics was performed using chip-based nanospray with an Advion TriVersa Nanomate (Advion) interfaced to the Thermo Exactive Orbitrap (Thermo Scientific). Briefy, a mass acquisition window from 200 to 2000 m/z and acquisition in positive and negative modes were used with a voltage of 1.2kV in positive mode and −1.5 kV in negative mode and an acquisition time of 72 s. Raw spectral data were processed as previously described^113^. Raw data were then converted to .mzXML (usingMSconvert^114^ with peakpick level 1), parsed with R and 50 spectra per sample (scan from 20 to 70) were averaged using XCMS42, with a signal cutoff at 2000. Te fles were aligned using the XCMS^115, 116^ grouping function using “mzClust” with a m/z-window of 22 ppm and a minimum coverage of 60%. Compound annotation was automated using both an exact mass search in compound libraries as well as applying the referenced Kendrick mass defect approach. Signal normalisation was performed by summing the intensities of all detected metabolites to a fixed value to produce a correction factor for the efficiency of ionisation. Exact masses were fitted to the lipid species library and subsequently annotated to the peak as described before^85^.

### Plasma proteomics

#### Sample preparation

Plasma was precleared by centrifugation at 3,000 g for 10 minutes and bound to 100 µL of calcium silicate matrix (CSM, 4 mg/mL) by rotation for 1 hour. The sample was centrifuged at 14,000 g for 1 minute and the supernatant was removed for further analysis. The pellet was washed in ammonium bicarbonate (50 mMoL, 1 mL) 3 times using the same centrifugation settings. The sample was then reduced for 30 minutes at 65°C using 200 µL of DL-dithiothreitol (DTT) premix (ADC 2%: ammonium bicarbonate 50 mMoL: DTT 1 MoL in the ratio of 50:49:1) and alkylated for 30 minutes in the dark with iodoacetamide (IAA) at 20 mMoL. Ammonium bicarbonate was added to dilute the ADC to 0.5%. Trypsin was added in the ratio of 1:25 trypsin to plasma and incubated overnight at 37°C. The ADC was precipitated with 1% formic acid (FA) and centrifuged at 14,000 g for 10 minutes. The peptides were isolated using solid phase EMPORE C18 discs which had been washed with 1 stem of methanol and 3 stem of 0.1% FA. The sample was left to bind to the column for 30 minutes before washing with 0.1% FA and eluting with 60% acetonitrile (ACN) with 0.1% FA and then 80% ACN with 0.1% FA. The ACN was removed by speed vacuum for 1 hour 15 minutes and freeze dried overnight. Peptide suspended in 30 µL of 0.1% FA and a peptide assay was performed to calculate the amount of peptides. 10 µL of peptides were removed from each sample and 0.1% FA added to equalise the volume and spiked with an internal standard protein (yeast alcohol dehydrogenase, ADH), with a known amount of 50 fmol injected for each run.

#### Waters NanoAcquity UPLC and Synapt G2S

Sample separation was performed using an Acquity UPLC Symmetry C18 trapping column (180 µm x 20mm, 5 µm) to remove salt and other impurities and a HSS T3 analytical column (75µm x 150mm, 1.8µm). Solvent A was compromised on 0.1% FA in HPLC grade water and solvent B contained 0.1% FA in ACN.

| Time<br>(minute) | Flow rate<br>($\mu\text{L}/\text{minute}$) | Solvent A<br>(Water + 0.1% FA) | Solvent B<br>(ACN + 0.1% FA) |
| --- | --- | --- | --- |
| 3 | 0.3 | 97 | 3 |
| 20 | 0.3 | 86 | 14 |
| 30 | 0.3 | 80 | 20 |
| 40 | 0.3 | 75 | 25 |
| 51-52.2 | 0.3 | 69 | 31 |
| 53-53.1 | 0.3 | 65 | 35 |
| 54 | 0.3 | 63 | 37 |
| 55 | 0.3 | 58 | 42 |
| 63 | 0.3 | 31 | 69 |
| 65 | 0.3 | 97 | 3 |
| 80 | 0.3 | 50 | 50 |
| 80.5 | 0.3 | 10 | 90 |
| 82.2-87.5 | 0.3 | 97 | 3 |
| 99.5 | 0.3 | 50 | 50 |
| 101.5 | 0.3 | 10 | 90 |
| 103.5-110 | 0.3 | 97 | 3 |

Table above shows the gradient in 110 minutes of solvent A and B used in LC ESI-MS/MS analysis. The flow rate of solvents was 0.3 µL/minute. Coupled directly to the Nano Acquity UPLC was a Water Synapt G2S mass spectrometer (Waters Corporation, Manchester, UK). The Synapt G2S includes a nano electrospray ionisation (ESI), StepWave ion guide, Quadrupole, TriWave and TOF (Supplementary Figure 2).

#### Proteomic data processing and analysis

Progenesis QI for Proteomics (Nonlinear Dynamics, Waters Corporation, UK) was employed to identify and quantify proteins. The human database from UniProtKB was downloaded and used in FASTA format. The proteomic raw data was searched using strict trypsin cleavage rules with a maximum of two missed cleavages. Cysteine (Carbamidomethyl C) was set as a fixed modification. Deamination N, Oxidation M and Phosphoryl STY were selected as variable modifications. Minimum of 2 fragments per peptide, minimum of 5 fragments per protein and minimum of 2 peptides per protein were set for parameters of identification. The maximum protein mass was set to 1000 kDa. The false rate discovery (FDR) for protein identification was set at a maximum rate of 1%. Then, proteomic data generated from using the Progenesis QI was exported to Microsoft Excel for further data analysis.

For differential analysis, we used LIMMA (v.3.37.4)^117^. Because we compared obese and post surgery patients, we performed a paired analysis. We then applied a threshold of 0.1% on ordinary qvalue.

To define whole blood specific genes, we exported GTEx project^118^ expression table (in TPMs), converted it into SummarizedExperiment container using SummarizedExperiment R package ((v.1.11.6); Morgan M, Obenchain V, Hester J, Pagès H SummarizedExperiment: SummarizedExperiment container. (2019)) and used teGeneRetrieval function from the TissueEnrich R package (v.1.2.1)^119^. This package relies on Human Protein Atlas^120^ to grouped genes as follows: Tissue Enriched (Genes with an expression level greater than 1 TPM that also have at least five-fold higher expression levels in a particular tissue compared to all other tissues), Group Enriched (Genes with an expression level greater than 1 TPM that also have at least five-fold higher expression levels in a group of 2-7 tissues compared to all other tissues, and that are not considered Tissue Enriched) and tissue Enhanced (Genes with an expression level greater than 1 TPM that also have at least five-fold higher expression levels in a particular tissue compared to the average levels in all other tissues, and that are not considered Tissue Enriched or Group Enriched). With default parameters, we identified 693 whole blood specific genes. Finally we interesected genes coding for differentially abundant proteins and whole blood specific genes.

#### Weighted correlation network analysis (WGCNA)

WGCNA^48^ is a correlation-based method that describes and visualizes networks of data points, whether they are gene expression estimates, metabolite concentrations or other phenotypic data. To increase statistical power, we merged the patient groups under the assumption that they share similar associations of metabolites and phenotypic traits. We followed the protocols of WGCNA to create metabolic networks. Metabolites are clustered into co-abundant "modules". Low correlations can be suppressed either in a continuous ("soft") manner or the discontinuous ("hard") thresholding used in constructing unweighted networks. To maintain scale-free topology, we estimated an applied power by computing soft-threshold with pickSoftThreshold function from WGCNA R package (v.1.64-1) ^121^. To build network, we used blockwiseModules function with the following options: TOMType = "signed", minModuleSize = 20, reassignThreshold = 0, mergeCutHeight = 0.25 and corType="bicor". Each obtained module is notated by a unique color. Additionally, we assigned a name to each consensus module. Each module abundance profile can be summarised by one representative metabolite: the module eigen metabolite. Specifically, the module eigen metabolite was defined as the first right-singular vector of the standardized module expression data^122^. We performed 3 analysis: extreme phenotypes (obese individuals and lipodystrophy patients were combined to get minimal sample size for network analysis), donors (all BD individuals) and a consensus analysis. We identified 8, 22 and 16 modules with donors, patients and consensus data respectively. Regarding consensus analysis, we considered 988 metabolites, of these, 375 were assigned to 15 different modules and the remaining 613 were put in an ad hoc extra module because they did not show any correlation. We computed eigenmodule and biochemical parameters correlations (leptin-adiponectin ratio (LAR), glucose (GLC), triglycerides (TG), total cholesterol (TC), high density lipoprotein (HDL-C), low-density lipoprotein (LDL-C), alanine amino-transferase (ALT), aspartate amino-transferase (AST), Homeostatic Model Assessment for Insulin Resistance (HOMA-IR) and adipose tissue insulin resistance (AT-IR) indexes and high-sensitivity C-reactive Protein (hsCRP) and also weight (WGT), BMI and age) using cor function from stats R base package (R version 3.5.0) and pearson method (default). P Value of each correlation was computed using corPvalueStudent function from WGCNA R package.

Pathways enrichment analysis were performed with MetaboAnalyst^123^ and in particular Pathway analysis module by submitting combined list HMDB identifiers for clusters C1 and C2, hyper-geometric test, relative-betweenness centrality topology analysis and KEGG database. In addition, we submitted these lists to the Reactome database.

### Multi-omic integration

#### Training datasets

We identified 16 BD individuals as lean, according to the following criteria: BMI < 25, glycaemia (GLUC) <5.4 mmol/L, TG <1.7 mmol/L, LDL <2.59 mmol/L, HDL >1 mmol/L for men and >1.3 mmol/L for women, HOMA score< 2.2. For training the multi-omics predictive model (see below), we used a reduced training dataset comprising the subset of individuals having measurements across all omics layers. This reduced set comprised 6 lean, 6 obese individuals and 10 lipodystrophy patients. For the clinical data, we first used multiple imputation by chained equations, as implemented in the mice R package (with default options) to impute missing values before construction of the training dataset. We used the same method to impute missing clinical values in the NASH cohort.

#### Variable selection: multivariable regression approach

For each of the omics layers considered independently, we used elastic-net penalised logistic regression as implemented in the glmnet R package to identify putative signatures that discriminated between all patients (i.e., lipodystrophy + obese) versus lean. We adjusted for age and sex by including them as unpenalised covariates in the multivariable model. Briefly, denoting by *M* the number of data layers, *N* the number of patients, and *X_m_* each layer’s training data matrix, where *m* = 1, …, *M*, we have *X_m_* ∈ ℝ^*N* × *P_m_*^ ordered such that corresponding rows in each matrix *X_m_* correspond to the same individual. For each individual, we also have a binary response, *y_i_*, and a vector ***W****_i_* comprising the covariates for which we wish to adjust; here, age and sex. We define the binary vector ***y*** = [*y*_1_, …, *y_N_*] and the *N* × 2 matrix *W* = [***W***_1_, …, ***W****_N_*]. For each layer, we perform an elastic net penalised logistic regression; i.e., for each *X_m_* we fit

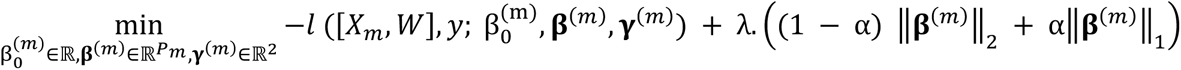

where: *l*(⋅) denotes the usual logistic regression likelihood function in which the (unpenalised) coefficients associated with age and sex are denoted by ***γ***^(*m*)^, the intercept term is denoted by 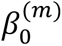, and the remaining regression coefficients are denoted by ***β***^(*m*)^; ‖***β***^(*m*)^‖_1_ and ‖***β***^(*m*)^‖_2_ represent the *l*_1_ and *l*_2_ norms respectively; *α* is fixed at 0.1; and the penalty parameter λ is selected via cross-validation. Since different cross-validation splits resulted in different choices for λ, we performed multiple rounds of cross-validation, and used the value of λ that resulted in the maximum number of selections. For each *X*_m_, we obtained a set of selected variables as those for which the estimated regression coefficients were non-zero.

#### Single layer predictive models

For each of the omics layers considered independently, we used the models fitted in the previous step to make predictions for the 96 individuals for which we had measurements across all omics layers.

#### Clinical predictive model

We trained a ridge-penalised logistic regression model predictive of the binary response (i.e. patient/lean status) using the clinical training dataset.

#### Multi-omics predictive model

We used all omic variables selected by the multivariable approach described above (i.e. the full collection of selected variables, across all data layers), together with the clinical covariates, to train a ridge-penalised logistic regression model predictive of the binary response (i.e. patient/lean status). We fitted this model using the reduced training dataset described above. We used this model to make predictions for the 96 individuals for which we had measurements across all omics layers. To allow us to make predictions for those individuals for which we only had measurements on a subset of the omics datasets, we additionally fitted models to each combination of subsets. A detailed analysis of this approach for selecting variables and training a multi-omic predictive model, including simulation studies to assess both predictive performance and ability to identify relevant predictors, is provided in Cabassi et al. (2020)^124^.

#### Validation of selected lipids

To further investigate the lipidomic signature, we prioritised a reduced set of 9 lipid species that had been selected into the signature. These 9 species satisfied the following criteria: (1) they were selected into the lipidomic signature; and (2) using the Mann-Whitney test with Storey’s q-value method to correct for multiple testing, we were able to reject the null hypothesis of no difference in distribution for these lipids in all of the following comparisons: (i) obese vs. lean; (ii) lipodystrophy vs. lean; and (iii) {obese and lipodystrophy} vs. lean. All tests were performed using data from the present study only. Of these 9 species, we were able to match 8 with lipid species that had been quantified in a subset of 1,507 participants of the Fenland study^83, 85^ which is a population-based cohort of 12,345 volunteers without diabetes born between 1950 and 1975 and recruited within the Cambridgeshire region between 2005 and 2015. We used linear regression analysis to test for association between plasma levels of the 8 lipid species selected into the lipidomic signature and all relevant CMS parameters quantified in both the reduced Fenland cohort, and the BD cohort, adjusting for age and sex, and using the Bonferroni method to control for multiple testing. To create a negative control set, we identified lipids that satisfied the following criteria: (1) they were not selected into the lipidomic signature; (2) they could be matched with lipid species that had been quantified in the reduced Fenland cohort; and (3) using the Mann-Whitney test with Storey’s q-value method to correct for multiple testing, we were unable to reject the null hypothesis of no difference in distribution for these lipids in any of the following comparisons: (i) obese vs. control; (ii) lipodystrophy vs. control; and (iii) {obese and lipodystrophy} vs. control. There were 37 lipid species that satisfied these criteria. We ranked these according to their mean absolute Pearson correlation with the 9 prioritised lipid species, and selected the 5 lowest ranking as our negative control set.

### Functional tests

#### Neutrophils Adhesion Method

Polymorphonuclear granulocytes were isolated via density gradient (1.078g/mL) from 3.2% sodium citrated whole blood within 2hours of venipuncture. Neutrophil purity was assessed by haematology analyser (Sysmex, XN-450) to ensure purity levels were satisfactory (≥90%) for subsequent functional assays. Isolated cells were incubated in a water bath at 37C for 30 minutes with fluorescently labelled Calcein-AM (4ug/mL, Molecular probes). Cells were washed twice with 1x PBS and resuspended at 2x106/ml in HEPES complete medium supplemented with calcium (1mM). 1.6x105 fluorescently labelled neutrophils were then added to relevant duplicate wells in a 96-well plate containing the following stimuli; fMLP, 10µM; DTT, 10mM; Pam3Cys, 20µg/ml; LBP+LPS, 50ng/mL and 20ng/mL; PAF, 1µM; PMA, 1µg/mL; TNF, 10ng/mL or HEPES only as a control in a final volume of in 100µl. Cells were incubated for 30 minutes at 37C in a 5% CO2 incubator, after which they were washed twice using 1x PBS before lysing in 100µl PBS with 0.5% triton. A 100% adhesion control was generated by lysing 1.6x105 fluorescently labelled neutrophils in 0.5% triton. Fluorescent intensity was measured using a Tecan Infinite® 200 PRO series plate reader (excitation of 485/20nm and emission of 535/25nm). The mean of duplicate values were calculated and the % adhesion over the hepes control calculated using the following formula: % adhesion = ((RFU stimuli – RFU HEPES)/ RFU 100% control) x 100.

CD63 Expression:

50ul of whole blood was incubated with antibodies:

|  |  |  |  |
| --- | --- | --- | --- |
| CD16 | PE | VEP13 | Miltenyi |
| CD63 | APC | H5C6 | Miltenyi |
| CD11b | APC | ICRF44 | BD Pharmingen™ |
| CD62L | FITC | Dreg 56 | BD Pharmingen™ |
| CD32 | FITC | FLI8.26 | BD Pharmingen™ |
| CD14 | APC | MφP9 | BD Pharmingen™ |

for 20 minutes, followed by a red cell lysis (BD FACS lyse) and resuspension in 0.2% formyl saline. Samples were analysed using flow cytometry (Beckman Coulter, FC500) within 4 hours. Neutrophils were identified using scatter properties and CD16 positivity. BD CompBeads were used to generate compensation controls. The median fluorescence intensity (MFI) for each surface marker was calculated using Kaluza Analysis Software (Beckman Coulter).

---

**Supplemental figure 1 - Related to Figure 1.**
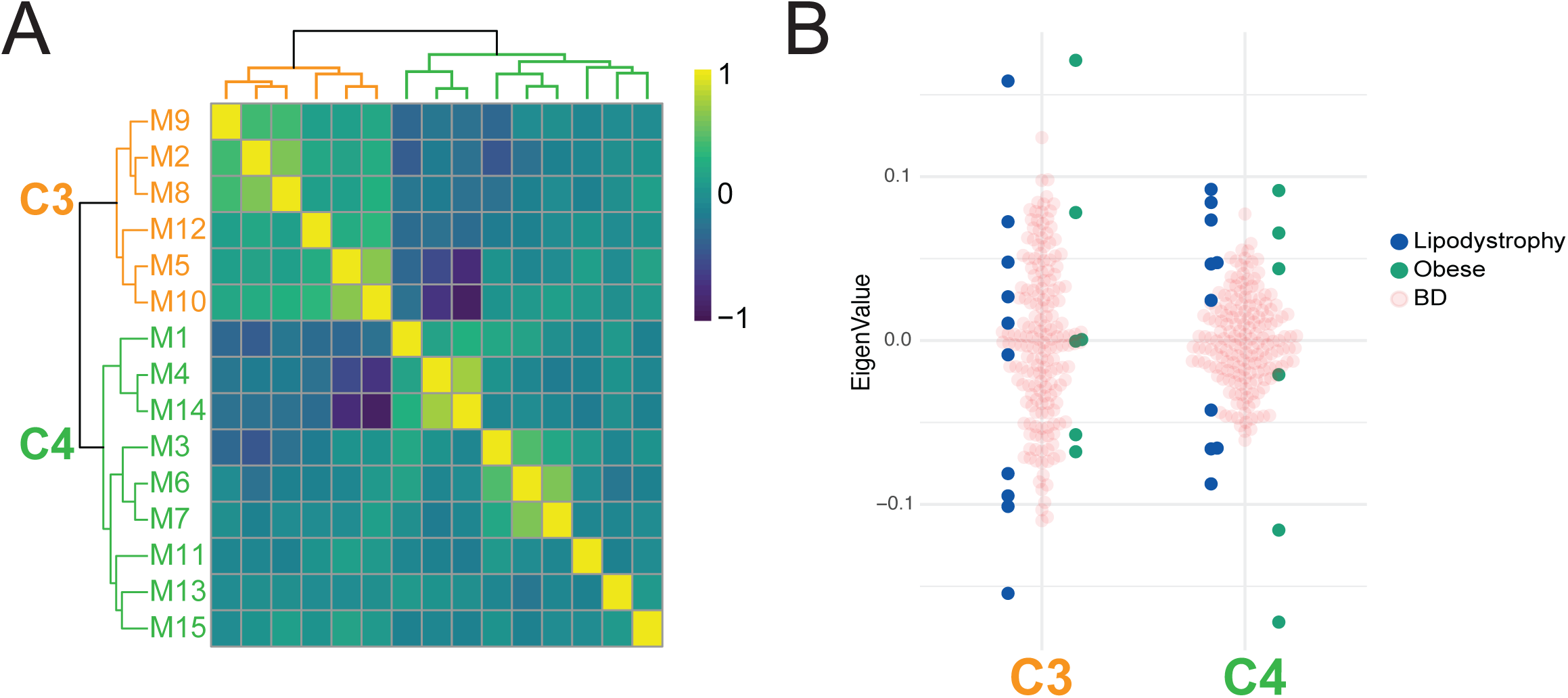
WGCNA analysis with BD individuals metabolite values and cluster functional annotation. A. Heatmap of BD individuals eigen-metabolites adjacencies in the consensus eigen-metabolites network. Each row and column correspond to one eigen-metabolite (labeled by consensus module color). The heat-map is color-coded by adjacency, yellow indicating high adjacency (positive correlation) and blue low adjacency (negative correlation) as shown by the color legend. B. Beeswarm plot using average eigen-metabolites per cluster. Colors indicate cohorts.

**Supplemental figure 2 - Related to Figure 2.**
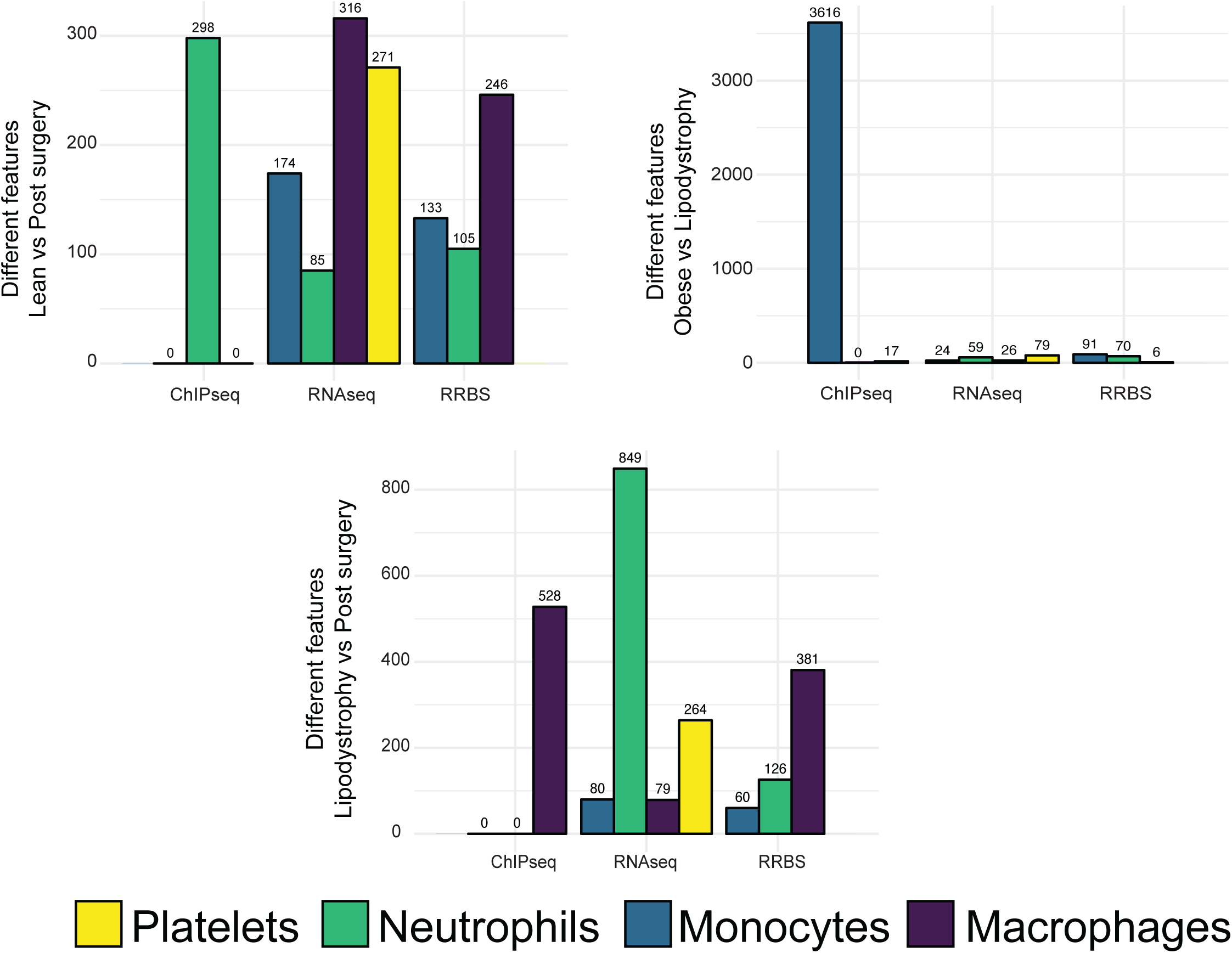
Summary plots of different feature numbers in all comparisons. Barplots showing the number of features significantly different for each comparison in H3K27ac distribution (ChIP-Seq), gene expression (RNA-Seq) and DNA methylation (RRBS). Each bar is color coded to represent the different cell types.

**Supplemental figure 3 - Related to Figure 3.**
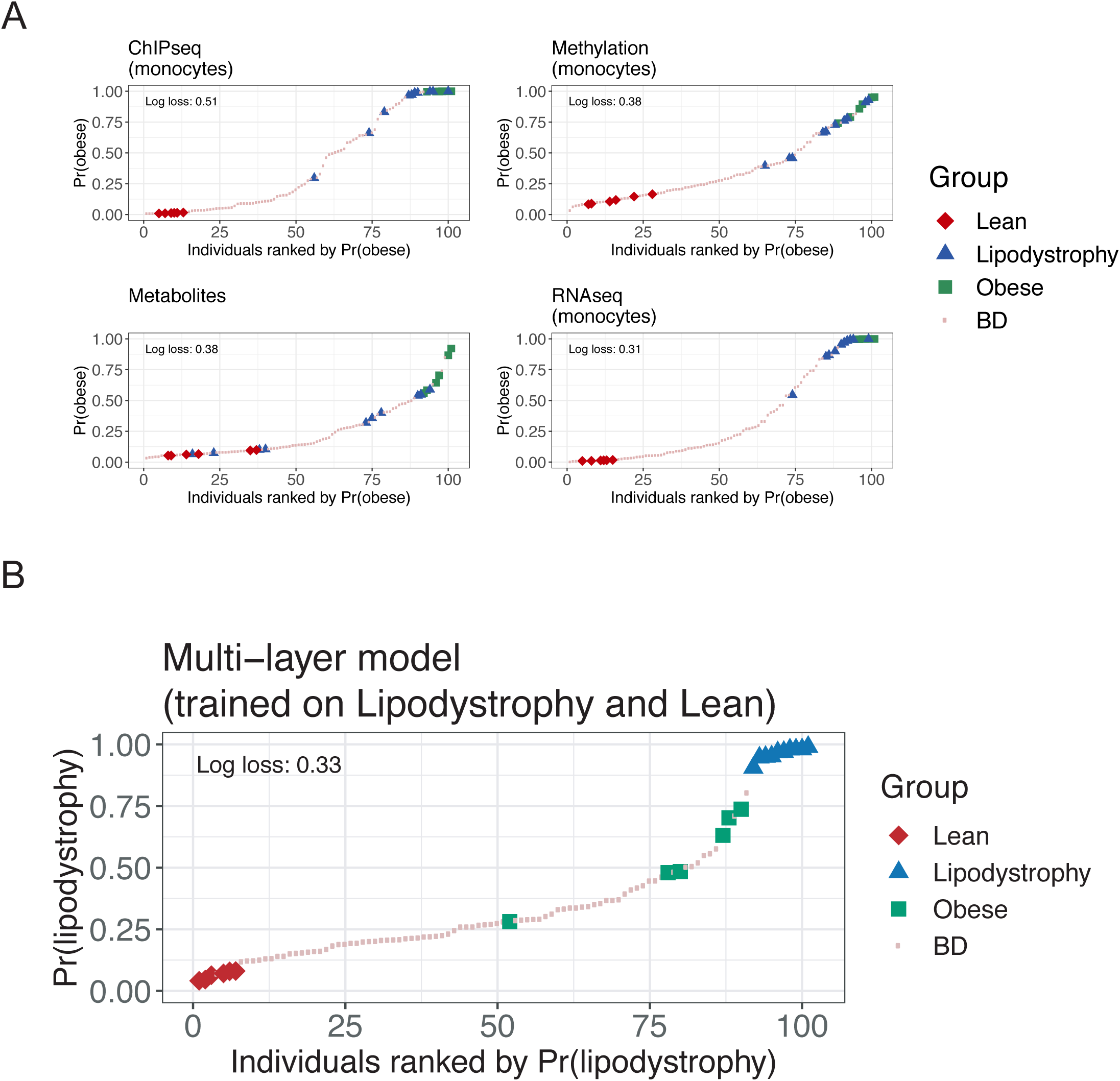
Multi-omic signatures of extreme phenotype groups and their use in prediction. A. Plots showing individuals ranked by their predicted probability of belonging to the obese group. As in Figure 3C, but for the Methylation (monocytes), RNA-Seq (monocytes), Metabolites, and ChIP-Seq (monocytes) data layers. B. Multi-omic model trained using lipodystrophy patients often predicts obese individuals to belong to the lipodystrophy group. As in Figure 3C (final plot), but training the multi-layer model using the Lipodystrophy and Lean groups (rather than the Obese and Lean groups). Using this model, Obese individuals were often predicted as belonging to the Lipodystrophy group.

**Supplemental figure 4 - Related to Figure 3.**
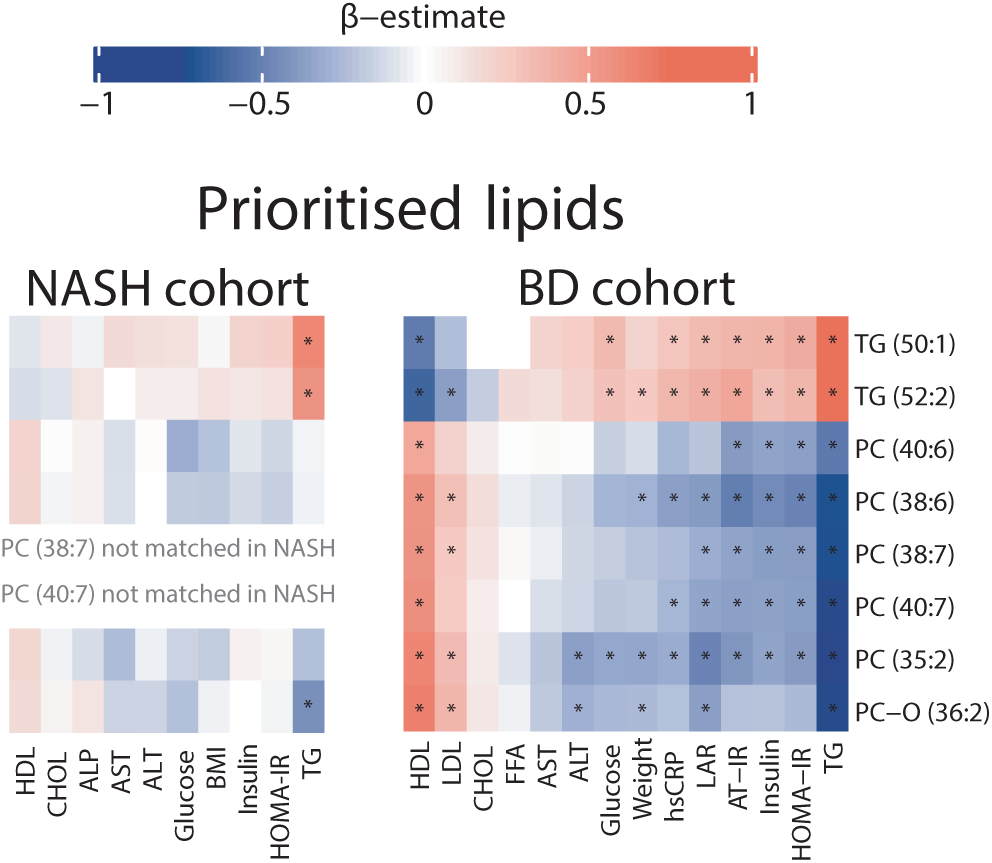
A common pattern of associations between the prioritised lipid species and known CMS risk factors. The pattern of association between the prioritised lipids and known CMS risk factors in the NASH cohort (NASH cohort; left) agrees with the results from the present study (BD cohort; right).

