## Extended Methods Metabolon for "Transcriptional, epigenetic and metabolic signatures in cardiometabolic syndrome defined by extreme phenotypes"

### mList$^{TM}$ REPORT

#### Biochemical profiling of rare disease cohort human plasma samples (BLUEPRINT)

**UCAM-0303B-15MLBL+**

AUTHOR: Edward D. Karoly, PhD

APPROVAL: Robert Mohney, PhD

DATE: May 06, 2016

Metabolon, Inc. • 617 Davis Drive, Suite 400, Durham, NC 27713 • (919) 572-1711 www.metabolon.com •
Contact:

#### Table of Contents

**Objective**

**Experimental Procedures**

**Results and Biological Interpretation**

*Metabolite Summary and Significantly Altered Biochemicals*

**Study Parameters**

*Data Quality: Instrument and Process Variability*

**Appendix**

*Metabolon Platform*

### Objective

#### *Purpose of Experiment*

The goal of this study was to biochemically profile human EDTA plasma samples submitted for analysis. These samples are associated with BLUEPRINT and includes samples (UCAM-09597 through UCAM-09819) originally submitted as part of UCAM-0303-15MLBL+.

### Experimental Procedures

#### *Experimental design*

Metabolon received 223 human EDTA plasma samples on October 7, 2015. The total number of samples submitted for analysis under this study is also presented below.

| Matrix | n |
|---|---|
| Human Plasma | 223 |

#### Results and Biological Interpretation

##### *Metabolite Summary and Significantly Altered Biochemicals*

The present dataset comprises a total of 1368 biochemicals, 909 compounds of known identity (named biochemicals) and 459 compounds of unknown structural identity (unnamed biochemicals). Information related to biochemical pathways, chemical properties, and public databases are included in the electronic data files that accompany this report.

| Matrix | Named Biochemicals | Unnamed Biochemicals | Total Biochemicals |
|---|---|---|---|
| Human Plasma | 909 | 459 | 1368 |

*[Biological interpretation is available as a component of the mView deliverable]*

#### Study Parameters

##### *Data Quality: Instrument and Process Variability*

| ***QC Sample*** | ***Measurement*** | ***Median RSD*** |
|---|---|---|
| Internal Standards | Instrument Variability | 4 % |
| Endogenous Biochemicals | Total Process Variability | 8 % |

Instrument variability was determined by calculating the median relative standard deviation (RSD) for the internal standards that were added to each sample prior to injection into the mass spectrometers. Overall process variability was determined by calculating the median RSD for all endogenous metabolites (i.e., non-instrument standards) present in 100% of the Client Matrix samples, which are technical replicates of pooled client samples. Values for instrument and process variability meet Metabolon's acceptance criteria as shown in the table above.

### Appendix

#### *Metabolon Platform*

**Sample Accessioning:** Following receipt, samples were inventoried and immediately stored at -80°C. Each sample received was accessioned into the Metabolon LIMS system and was assigned by the LIMS a unique identifier that was associated with the original source identifier only. This identifier was used to track all sample handling, tasks, results, etc. The samples (and all derived aliquots) were tracked by the LIMS system. All portions of any sample were automatically assigned their own unique identifiers by the LIMS when a new task was created; the relationship of these samples was also tracked. All samples were maintained at -80°C until processed.

**Sample Preparation:** Samples were prepared using the automated MicroLab STAR® system from Hamilton Company. Several recovery standards were added prior to the first step in the extraction process for QC purposes. To remove protein, dissociate small molecules bound to protein or trapped in the precipitated protein matrix, and to recover chemically diverse metabolites, proteins were precipitated with methanol under vigorous shaking for 2 min (Glen Mills GenoGrinder 2000) followed by centrifugation. The resulting extract was divided into five fractions: two for analysis by two separate reverse phase (RP)/UPLC-MS/MS methods with positive ion mode electrospray ionization (ESI), one for analysis by RP/UPLC-MS/MS with negative ion mode ESI, one for analysis by HILIC/UPLC-MS/MS with negative ion mode ESI, and one sample was reserved for backup. Samples were placed briefly on a TurboVap® (Zymark) to remove the organic solvent. The sample extracts were stored overnight under nitrogen before preparation for analysis.

**QA/QC:** Several types of controls were analyzed in concert with the experimental samples: a pooled matrix sample generated by taking a small volume of each experimental sample (or alternatively, use of a pool of well-characterized human plasma) served as a technical replicate throughout the data set; extracted water samples served as process blanks; and a cocktail of QC standards that were carefully chosen not to interfere with the measurement of endogenous compounds were spiked into every analyzed sample, allowed instrument performance monitoring and aided chromatographic alignment. Tables 1 and 2 describe these QC samples and standards. Instrument variability was determined by calculating the median relative standard deviation (RSD) for the standards that were added to each sample prior to injection into the mass spectrometers. Overall process variability was determined by calculating the median RSD for all endogenous metabolites (i.e., non-instrument standards) present in 100% of the pooled matrix samples. Experimental samples were randomized across the platform run with QC samples spaced evenly among the injections, as outlined in Figure 1.

**Table 1: Description of Metabolon QC Samples**

| Type | Description | Purpose |
|---|---|---|
| MTRX | Large pool of human plasma maintained by Metabolon that has been characterized extensively. | Assure that all aspects of the Metabolon process are operating within specifications. |
| CMTRX | Pool created by taking a small aliquot from every customer sample. | Assess the effect of a non-plasma matrix on the Metabolon process and distinguish biological variability from process variability. |
| PRCS | Aliquot of ultra-pure water | Process Blank used to assess the contribution to compound signals from the process. |
| SOLV | Aliquot of solvents used in extraction. | Solvent Blank used to segregate contamination sources in the extraction. |

**Table 2: Metabolon QC Standards**

| Type | Description | Purpose |
|---|---|---|
| RS | Recovery Standard | Assess variability and verify performance of extraction and instrumentation. |
| IS | Internal Standard | Assess variability and performance of instrument. |

**Figure 1. Preparation of client-specific technical replicates.** A small aliquot of each client sample (colored cylinders) is pooled to create a CMTRX technical replicate sample (multi-colored cylinder), which is then injected periodically throughout the platform run. Variability among consistently detected biochemicals can be used to calculate an estimate of overall process and platform variability.

**Ultrahigh Performance Liquid Chromatography-Tandem Mass Spectroscopy (UPLC-MS/MS):** All methods utilized a Waters ACQUITY ultra-performance liquid chromatography (UPLC) and a Thermo Scientific Q-Exactive high resolution/accurate mass spectrometer interfaced with a heated electrospray ionization (HESI-II) source and Orbitrap mass analyzer operated at 35,000 mass resolution. The sample extract was dried then reconstituted in solvents compatible to each of the four methods. Each reconstitution solvent contained a series of standards at fixed concentrations to ensure injection and chromatographic consistency. One aliquot was analyzed using acidic positive ion conditions, chromatographically optimized for more hydrophilic compounds. In this method, the extract was gradient eluted from a C18 column (Waters UPLC BEH C18-2.1x100 mm, 1.7 µm) using water and methanol, containing 0.05% perfluoropentanoic acid (PFPA) and 0.1% formic acid (FA). Another aliquot was also analyzed using acidic positive ion conditions, however it was chromatographically optimized for more hydrophobic compounds. In this method, the extract was gradient eluted from the same afore mentioned C18 column using methanol, acetonitrile, water, 0.05% PFPA and 0.01% FA and was operated at an overall higher organic content. Another aliquot was analyzed using basic negative ion optimized conditions using a separate dedicated C18 column. The basic extracts were gradient eluted from the column using methanol and water, however with 6.5mM Ammonium Bicarbonate at pH 8. The fourth aliquot was analyzed via negative ionization following elution from a HILIC column (Waters UPLC BEH Amide 2.1x150 mm, 1.7 µm) using a gradient consisting of water and acetonitrile with 10mM Ammonium Formate, pH 10.8. The MS analysis alternated between MS and data-dependent $MS^n$ scans using dynamic exclusion. The scan range varied slighted between methods but covered 70-1000 m/z. Raw data files are archived and extracted as described below.

**Bioinformatics:** The informatics system consisted of four major components, the Laboratory Information Management System (LIMS), the data extraction and peak-identification software, data processing tools for QC and compound identification, and a collection of information interpretation and visualization tools for use by data analysts. The hardware and software foundations for these informatics components were the LAN backbone, and a database server running Oracle 10.2.0.1 Enterprise Edition.

**LIMS:** The purpose of the Metabolon LIMS system was to enable fully auditable laboratory automation through a secure, easy to use, and highly specialized system. The scope of the

Metabolon LIMS system encompasses sample accessioning, sample preparation and instrumental analysis and reporting and advanced data analysis. All of the subsequent software systems are grounded in the LIMS data structures. It has been modified to leverage and interface with the in-house information extraction and data visualization systems, as well as third party instrumentation and data analysis software.

**Data Extraction and Compound Identification:** Raw data was extracted, peak-identified and QC processed using Metabolon's hardware and software. These systems are built on a web-service platform utilizing Microsoft's .NET technologies, which run on high-performance application servers and fiber-channel storage arrays in clusters to provide active failover and load-balancing. Compounds were identified by comparison to library entries of purified standards or recurrent unknown entities. Metabolon maintains a library based on authenticated standards that contains the retention time/index (RI), mass to charge ratio (*m/z)*, and chromatographic data (including MS/MS spectral data) on all molecules present in the library. Furthermore, biochemical identifications are based on three criteria: retention index within a narrow RI window of the proposed identification, accurate mass match to the library +/- 10 ppm, and the MS/MS forward and reverse scores between the experimental data and authentic standards. The MS/MS scores are based on a comparison of the ions present in the experimental spectrum to the ions present in the library spectrum. While there may be similarities between these molecules based on one of these factors, the use of all three data points can be utilized to distinguish and differentiate biochemicals. More than 3300 commercially available purified standard compounds have been acquired and registered into LIMS for analysis on all platforms for determination of their analytical characteristics. Additional mass spectral entries have been created for structurally unnamed biochemicals, which have been identified by virtue of their recurrent nature (both chromatographic and mass spectral). These compounds have the potential to be identified by future acquisition of a matching purified standard or by classical structural analysis.

**Curation:** A variety of curation procedures were carried out to ensure that a high quality data set was made available for statistical analysis and data interpretation. The QC and curation processes were designed to ensure accurate and consistent identification of true chemical entities, and to remove those representing system artifacts, mis-assignments, and background noise. Metabolon data analysts use proprietary visualization and interpretation software to confirm the consistency of peak identification among the various samples. Library matches for each compound were checked for each sample and corrected if necessary.

**Metabolite Quantification and Data Normalization:** Peaks were quantified using area-under-the-curve. For studies spanning multiple days, a data normalization step was performed to correct variation resulting from instrument inter-day tuning differences. Essentially, each compound was corrected in run-day blocks by registering the medians to equal one (1.00) and normalizing each data point proportionately (termed the "block correction"; Figure 2). For studies that did not require more than one day of analysis, no normalization is necessary, other than for purposes of data visualization. In certain instances, biochemical data may have been normalized to an additional factor (e.g., cell counts, total protein as determined by Bradford assay, osmolality, etc.) to account for differences in metabolite levels due to differences in the amount of material present in each sample.

**Figure 2: Visualization of data normalization steps for a multiday platform run.**

### mList+™ FINAL REPORT

#### Biochemical profiling of BLUEPRINT plasma samples

##### UCAM-0312D-15MLBL+

CLIENT: Willem Ouwehand PhD

AUTHOR: Gregory Michelotti, PhD

APPROVAL: Bryan Wittmann, PhD

DATE: August 2, 2016

Metabolon, Inc. • 617 Davis Drive, Suite 400, Durham, NC 27713 • (919) 572-1711 www.metabolon.com •
Contact:

Table of Contents

**Objective**

**Experimental Procedures**

**Results and Biological Interpretation**

*[Biological Interpretation is available as a component of mView service deliverable.]*

***Metabolite Summary and Significantly Altered Biochemicals***

**Study Parameters**

***Data Quality: Instrument and Process Variability***

**Appendix**

***Metabolon Platform***

***Statistical Methods and Terminology***

#### Objective

The goal of the study was to identify metabolic differences in rare disease group samples that are associated with BLUEPRINT samples from the Ouwehand group. As per delineated in a client-provided manifest, 20 plasma samples from the current study were merged with 223 samples from the previous 0303B study, totaling 243 samples.

#### Experimental Procedures

##### *Experimental design*

Global biochemical profiles were determined in human plasma samples associated with samples below.

| Matrix | Sample n | Description |
|---|---|---|
| Human Plasma | 243 | Willem Ouwehand's BLUEPRINT samples |

#### Results and Biological Interpretation

*[Biological Interpretation is available as a component of mView service deliverable.]*

##### *Metabolite Summary and Significantly Altered Biochemicals*

Numbers of compounds of known structural identity (named biochemicals) as well as compounds of unknown structural identity (unnamed biochemicals) detected in the present datasets are presented in the table below. Information related to biochemical pathways, chemical properties, and public databases are included in the electronic data files that accompany this report.

| ***Matrix*** | ***Named Biochemicals*** | ***Unnamed Biochemicals*** | ***Total Biochemicals*** |
|---|---|---|---|
| Human Plasma | 793 | 362 | 1155 |

#### Study Parameters

##### *Data Quality: Instrument and Process Variability*

| *QC Sample* | *Measurement* | *Median RSD* |
|---|---|---|
| Internal Standards | Instrument Variability | 4% |
| Endogenous Biochemicals | Total Process Variability | 9% |

Instrument variability was determined by calculating the median relative standard deviation (RSD) for the internal standards that were added to each sample prior to injection into the mass spectrometers. Overall process variability was determined by calculating the median RSD for all endogenous metabolites (i.e., non-instrument standards) present in 100% of the Client Matrix samples, which are technical replicates of pooled client samples. Values for instrument and process variability as shown in the table above meet Metabolon's acceptance criteria.

#### Appendix

##### *Metabolon Platform*

**Sample Accessioning:** Following receipt, samples were inventoried and immediately stored at -80$^{o}$C. Each sample received was accessioned into the Metabolon LIMS system and was assigned by the LIMS a unique identifier that was associated with the original source identifier only. This identifier was used to track all sample handling, tasks, results, etc. The samples (and all derived aliquots) were tracked by the LIMS system. All portions of any sample were automatically assigned their own unique identifiers by the LIMS when a new task was created; the relationship of these samples was also tracked. All samples were maintained at -80$^{o}$C until processed.

**Sample Preparation:** Samples were prepared using the automated MicroLab STAR® system from Hamilton Company. Several recovery standards were added prior to the first step in the extraction process for QC purposes. To remove protein, dissociate small molecules bound to protein or trapped in the precipitated protein matrix, and to recover chemically diverse metabolites, proteins were precipitated with methanol under vigorous shaking for 2 min (Glen Mills GenoGrinder 2000) followed by centrifugation. The resulting extract was divided into five fractions: two for analysis by two separate reverse phase (RP)/UPLC-MS/MS methods with positive ion mode electrospray ionization (ESI), one for analysis by RP/UPLC-MS/MS with negative ion mode ESI, one for analysis by HILIC/UPLC-MS/MS with negative ion mode ESI, and one sample was reserved for backup. Samples were placed briefly on a TurboVap® (Zymark) to remove the organic solvent. The sample extracts were stored overnight under nitrogen before preparation for analysis.

**QA/QC:** Several types of controls were analyzed in concert with the experimental samples: a pooled matrix sample generated by taking a small volume of each experimental sample (or alternatively, use of a pool of well-characterized human plasma) served as a technical replicate throughout the data set; extracted water samples served as process blanks; and a cocktail of QC standards that were carefully chosen not to interfere with the measurement of endogenous compounds were spiked into every analyzed sample, allowed instrument performance monitoring and aided chromatographic alignment. Tables 1 and 2 describe these QC samples and standards. Instrument variability was determined by calculating the median relative standard deviation (RSD) for the standards that were added to each sample prior to injection into the mass spectrometers. Overall process variability was determined by calculating the median RSD for all endogenous metabolites (i.e., non-instrument standards) present in 100% of the pooled matrix samples. Experimental samples were randomized across the platform run with QC samples spaced evenly among the injections, as outlined in Figure 1.

**Table 1: Description of Metabolon QC Samples**

| Type | Description | Purpose |
|---|---|---|
| MTRX | Large pool of human plasma maintained by Metabolon that has been characterized extensively. | Assure that all aspects of the Metabolon process are operating within specifications. |
| CMTRX | Pool created by taking a small aliquot from every customer sample. | Assess the effect of a non-plasma matrix on the Metabolon process and distinguish biological variability from process variability. |
| PRCS | Aliquot of ultra-pure water | Process Blank used to assess the contribution to compound signals from the process. |
| SOLV | Aliquot of solvents used in extraction. | Solvent Blank used to segregate contamination sources in the extraction. |

**Table 2: Metabolon QC Standards**

| Type | Description | Purpose |
|---|---|---|
| RS | Recovery Standard | Assess variability and verify performance of extraction and instrumentation. |
| IS | Internal Standard | Assess variability and performance of instrument. |

**Figure 1. Preparation of client-specific technical replicates.** A small aliquot of each client sample (colored cylinders) is pooled to create a CMTRX technical replicate sample (multi-colored cylinder), which is then injected periodically throughout the platform run. Variability among consistently detected biochemicals can be used to calculate an estimate of overall process and platform variability.

**Ultrahigh Performance Liquid Chromatography-Tandem Mass Spectroscopy (UPLC-MS/MS):** All methods utilized a Waters ACQUITY ultra-performance liquid chromatography (UPLC) and a Thermo Scientific Q-Exactive high resolution/accurate mass spectrometer interfaced with a heated electrospray ionization (HESI-II) source and Orbitrap mass analyzer operated at 35,000 mass resolution. The sample extract was dried then reconstituted in solvents compatible to each of the four methods. Each reconstitution solvent contained a series of standards at fixed concentrations to ensure injection and chromatographic consistency. One aliquot was analyzed using acidic positive ion conditions, chromatographically optimized for more hydrophilic compounds. In this method, the extract was gradient eluted from a C18 column (Waters UPLC BEH C18-2.1x100 mm, 1.7 µm) using water and methanol, containing 0.05% perfluoropentanoic acid (PFPA) and 0.1% formic acid (FA). Another aliquot was also analyzed using acidic positive ion conditions, however it was chromatographically optimized for more hydrophobic compounds. In this method, the extract was gradient eluted from the same afore mentioned C18 column using methanol, acetonitrile, water, 0.05% PFPA and 0.01% FA and was operated at an overall higher organic content. Another aliquot was analyzed using basic negative ion optimized conditions using a separate dedicated C18 column. The basic extracts were gradient eluted from the column using methanol and water, however with 6.5mM Ammonium Bicarbonate at pH 8. The fourth aliquot was analyzed via negative ionization following elution from a HILIC column (Waters UPLC BEH Amide 2.1x150 mm, 1.7 µm) using a gradient consisting of water and acetonitrile with 10mM Ammonium Formate, pH 10.8. The MS analysis alternated between MS and data-dependent $MS^n$ scans using dynamic exclusion. The scan range varied slighted between methods but covered 70-1000 m/z. Raw data files are archived and extracted as described below.

**Bioinformatics:** The informatics system consisted of four major components, the Laboratory Information Management System (LIMS), the data extraction and peak-identification software, data processing tools for QC and compound identification, and a collection of information interpretation and visualization tools for use by data analysts. The hardware and software foundations for these informatics components were the LAN backbone, and a database server running Oracle 10.2.0.1 Enterprise Edition.

**LIMS:** The purpose of the Metabolon LIMS system was to enable fully auditable laboratory automation through a secure, easy to use, and highly specialized system. The scope of the Metabolon LIMS system encompasses sample accessioning, sample preparation and instrumental analysis and reporting and advanced data analysis. All of the subsequent software systems are grounded in the LIMS data structures. It has been modified to leverage and interface with the in-house information extraction and data visualization systems, as well as third party instrumentation and data analysis software.

**Data Extraction and Compound Identification:** Raw data was extracted, peak-identified and QC processed using Metabolon's hardware and software. These systems are built on a web-service platform utilizing Microsoft's .NET technologies, which run on high-performance application servers and fiber-channel storage arrays in clusters to provide active failover and load-balancing. Compounds were identified by comparison to library entries of purified standards or recurrent unknown entities. Metabolon maintains a library based on authenticated standards that contains the retention time/index (RI), mass to charge ratio (*m/z)*, and chromatographic data (including MS/MS spectral data) on all molecules present in the library. Furthermore, biochemical identifications are based on three criteria: retention index within a narrow RI window of the proposed identification, accurate mass match to the library +/- 10 ppm, and the MS/MS forward and reverse scores between the experimental data and authentic standards. The MS/MS scores are based on a comparison of the ions present in the experimental spectrum to the ions present in the library spectrum. While there may be similarities between these molecules based on one of these factors, the use of all three data points can be utilized to distinguish and differentiate biochemicals. More than 3300 commercially available purified standard compounds have been acquired and registered into LIMS for analysis on all platforms for determination of their analytical characteristics. Additional mass spectral entries have been created for structurally unnamed biochemicals, which have been identified by virtue of their recurrent nature (both chromatographic and mass spectral). These compounds have the potential to be identified by future acquisition of a matching purified standard or by classical structural analysis.

**Curation:** A variety of curation procedures were carried out to ensure that a high quality data set was made available for statistical analysis and data interpretation. The QC and curation processes were designed to ensure accurate and consistent identification of true chemical entities, and to remove those representing system artifacts, mis-assignments, and background noise. Metabolon data analysts use proprietary visualization and interpretation software to confirm the consistency of peak identification among the various samples. Library matches for each compound were checked for each sample and corrected if necessary.

**Metabolite Quantification and Data Normalization:** Peaks were quantified using area-under-the-curve. For studies spanning multiple days, a data normalization step was performed to correct variation resulting from instrument inter-day tuning differences. Essentially, each compound was corrected in run-day blocks by registering the medians to equal one (1.00) and normalizing each data point proportionately (termed the "block correction"; Figure 2). For studies that did not require more than one day of analysis, no normalization is necessary, other than for purposes of data visualization. In certain instances, biochemical data may have been normalized to an additional factor (e.g., cell counts, total protein as determined by Bradford assay, osmolality, etc.) to account for differences in metabolite levels due to differences in the amount of material present in each sample.

**Figure 2: Visualization of data normalization steps for a multiday platform run.**

#### *Statistical Methods and Terminology*

**Statistical Calculations:** For many studies, two types of statistical analysis are usually performed: (1) significance tests and (2) classification analysis. Standard statistical analyses are performed in ArrayStudio on log transformed data. For those analyses not standard in ArrayStudio, the programs R (http://cran.r-project.org/) or JMP are used. Below are examples of frequently employed significance tests and classification methods followed by a discussion of p- and q-value significance thresholds.

1. **Welch's two-sample *t*-test**

Welch's two-sample *t*-test is used to test whether two unknown means are different from two independent populations.

This version of the two-sample *t*-test allows for unequal variances (variance is the square of the standard deviation) and has an *approximate t*-distribution with degrees of freedom estimated using Satterthwaite's approximation. The test statistic is given by *t* , and the degrees of freedom is given by , where , are the sample means, $s_1$, $s_2$, are the sample standard deviations, and $n_1$, $n_2$ are the samples sizes from groups 1 and 2, respectively. We typically use a two-sided test (tests whether the means are different) as opposed to a one-sided test (tests whether one mean is greater than the other).

**2. Matched Pairs *t*-test**

The matched pairs *t*-test is used to test whether two unknown means are different from paired observations taken on the same subjects.

The matched pairs *t*-test is equivalent to the one-sample *t*-test performed on the differences of the observations taken on each subject (i.e., calculate ($x_1 - x_2$) for each subject; test whether the mean difference is zero or not). The test statistic is given by , with $n - 1$ degrees of freedom, where , are the sample means for groups 1 and 2, respectively, $s_d$ is the standard deviation of the differences, $n$ is the number of *subjects* (so there are $2n$ observations).

**3. One-way ANOVA**

ANOVA stands for analysis of variance. For ANOVA, it is assumed that all populations have the same variances. One-way ANOVA is used to test whether at least two unknown means are all equal or whether at least one pair of means is different. For the case of two means, ANOVA gives the same result as a two-sided *t*-test with a pooled estimate of the variance.

An ANOVA uses an F-test which has two parameters – the numerator degrees of freedom and the denominator degrees of freedom. The degrees of freedom in the numerator are equal to $g - 1$, where $g$ is the number of groups. If $n$ is the total number of observations ($n_1 + n_2$), then, the denominator degrees of freedom is equal to $n - g$. The F-statistic is the ratio of the between-groups variance to the within-groups variance, hence the higher the F-statistic the more evidence we have that the means are different.

Often within ANOVA, one performs linear contrasts for specific comparisons of interest. For example, suppose we have three groups A, B, C, then examples of some contrasts are A vs. B, the average of A and B vs. C, etc. For single-degree of freedom contrasts, these give the same result as a two-sided *t*-test with the pooled estimate of the variance from the ANOVA and degrees of freedom $n - g$. Below, we show the three formulas for A vs. B from a three group design as shown above. The numerator is same in each case, but the denominator differs by the estimates of the variances, and the degrees of freedom are different for each (if the theoretical assumptions hold, then the contrast has the most power, as it has the largest degrees of freedom).

Welch's two-sample *t*-test
By *t* , and the degrees of freedom is given by

Two-sample *t*-test with pooled estimate of variance from A and B
where , where the degrees of freedom is $n_A + n_B – 2$.
The contrast from the ANOVA,
where , where the degrees of freedom is given by where the degrees of freedom is $n_A + n_B + n_C – 3$.

#### 4. Two-way ANOVA

ANOVA stands for analysis of variance. For ANOVA, it is assumed that all populations have the same variances. For a two-way ANOVA, three statistical tests are typically performed: the main effect of each factor and the interaction. Suppose we have two factors A and B, where A represent the genotype and B represent the diet in a mouse study. Suppose each of these factors has two levels (A: wild type, knock out; B: standard diet, high fat diet). For this example, there are 4 combinations ("treatments"): A1B1, A1B2, A2B1, A2B2. The overall ANOVA F-test gives the p-value for testing whether all four of these means are equal or whether at least one pair is different. However, we are also interested in the effect of the genotype and diet. A main effect is a contrast that tests one factor across the levels of the other factor. Hence the A main effect compares (A1B1 + A1B2)/2 vs. (A2B1 + A2B2)/2, and the B-main effect compares (A1B1 + A2B2)/2 vs. (A1B2 + A2B2)/2. The interaction is a contrast that tests whether the mean difference for one factor depends on the level of the other factor, which is (A1B2 + A2B1)/2 vs. (A1B1 + A2B2)/2.

Some sample plots follow. For the first plot, there is a B main effect, but no A main effect and no interaction, as the effect of B does not depend on the level of A. For the second plot, notice how the mean difference for B is the same at each level of A and the difference in A is the same for each level of B, hence there is no statistical interaction. The final plot also has main effects for A and B, but here also has an interaction: we see the effect of B depends on the level of A (0 for A1 but 2 for A2), i.e., the effect of the diet depends on the genotype. We also see here the interpretation of the main effects depends on whether there is an interaction or not.

#### 5. Two-way Repeated Measures ANOVA

This is typically an ANOVA where one factor is applied to each subject and the second factor is a time point. See two-way ANOVA as many of the details are similar except that the model takes into account the repeated measures, i.e., the treatments are given to the same subject over time. The two main effects and the interaction are assessed, with particular interest to the interaction, as this shows where the time profiles are parallel or not for the treatments (parallel mean no interaction).

One additional note, the standard analysis assumes a condition referred to as compound symmetry, which assumes the correlation between each pair of levels of the repeated-measures factor is the same. Thus, for the case of time, it assumes the correlation is the same between time points 1 and 2, 1 and 3, and 2 and 3.

**6. Correlation**

Correlation measures the strength and direction of a *linear* association between two variables. The statistical test for correlation tests whether the true correlation is zero or not.

The square of the correlation is the percentage of the total variation explained by a linear relationship between the two variables. Thus, with large sample sizes there may be a sample correlation of 0.1 that is statistically significant. This means we have high confidence that the true correlation is zero, however, only 100*(0.1*0.1)% = 1% of the variation of one variable is explained by a linear relationship with the other variable, so while there is an association, it has little predictive ability.

**7. Hotelling's $T^2$ test**

The Hotelling's $T^2$ test is a multivariate generalization of the *t*-test, but here we are testing whether the mean vectors are different or not (the vector consists of multiple metabolites).

The Hotelling statistic is: , where $n_x$ and $n_y$ are the numbers of samples in each group, is the mean vector of the variables from group 1, is the mean vector of variables from group 2 and **S** is the pooled estimate of the variance-covariance matrix of the variables. This analysis assumes the underlying variance-covariance matrix is the same for each group. Notice that in the case of uncorrelated variables, this is simply a weighted average of the squared mean differences with weights inversely proportional to the sample variances (i.e., the metabolites less variable within a group are given higher weights).

**8. p-values**

For statistical significance testing, p-values are given. The lower the p-value, the more evidence we have that the null hypothesis (typically that two population means are equal) is not true. If "statistical significance" is declared for p-values less than

0.05, then 5% of the time we incorrectly conclude the means are different, when actually they are the same.

The p-value is the probability that the test statistic is at least as extreme as observed in this experiment given that the null hypothesis is true. Hence, the more extreme the statistic, the lower the p-value and the more evidence the data gives against the null hypothesis.

#### 9. q-values

The level of 0.05 is the false positive rate when there is one test. However, for a large number of tests we need to account for false positives. There are different methods to correct for multiple testing. The oldest methods are family-wise error rate adjustments (Bonferroni, Tukey, etc.), but these tend to be extremely conservative for a very large number of tests. With gene arrays, using the False Discovery Rate (FDR) is more common. The family-wise error rate adjustments give one a high degree of confidence that there are zero false discoveries. However, with FDR methods, one can allow for a small number of false discoveries. The FDR for a given set of compounds can be estimated using the q-value (see Storey J and Tibshirani R. (2003) Statistical significance for genomewide studies. Proc. Natl. Acad. Sci. USA 100: 9440-9445; PMID: 12883005).

In order to interpret the q-value, the data must first be sorted by the p-value then choose the cutoff for significance (typically $p<0.05$). The q-value gives the false discovery rate for the selected list (i.e., an estimate of the proportion of false discoveries for the list of compounds whose p-value is below the cutoff for significance). For Table 1 below, if the whole list is declared significant, then the false discovery rate is approximately 10%. If everything from Compound 079 and above is declared significant, then the false discovery rate is approximately 2.5%.

Table 1: Example of q-value interpretation

#### 10. Random Forest

Random forest is a supervised classification technique based on an ensemble of decision trees (see Breiman L. (2001) Random Forests. Machine Learning. 45: 5-32; http://link.springer.com/article/10.1023%2FA%3A1010933404324). For a given decision tree, a random subset of the data with identifying true class information is selected to build the tree ("bootstrap sample" or "training set"), and then the remaining data, the "out-of-bag" (OOB) variables, are passed down the tree to obtain a class prediction for each sample. This process is repeated thousands of times to produce the forest. The final classification of each sample is determined by computing the class prediction frequency ("votes") for the OOB variables over the whole forest. For example, suppose the random forest consists of 50,000 trees and that 25,000 trees had a prediction for sample 1. Of these 25,000, suppose 15,000 trees classified the sample as belonging to Group A and the remaining 10,000 classified it as belonging to Group B. Then the votes are 0.6 for Group A and 0.4 for Group B, and hence the final classification is Group A. This method is unbiased since the prediction for each sample is based on trees built from a subset of samples that do not include that sample. When the full forest is grown, the class predictions are compared to the true classes, generating the "OOB error rate" as a measure of prediction accuracy. Thus, the prediction accuracy is an unbiased estimate of how well one can predict sample class in a new data set. Random forest has several advantages – it makes no parametric assumptions, variable selection is not needed, it does not overfit, it is invariant to transformation, and it is fairly easy to implement with R.

To determine which variables (biochemicals) make the largest contribution to the classification, a "variable importance" measure is computed. We use the "Mean Decrease Accuracy" (MDA) as this metric. The MDA is determined by randomly permuting a variable, running the observed values through the trees, and then reassessing the prediction accuracy. If a variable is not important, then this procedure will have little change in the accuracy of the class prediction (permuting random noise will give random noise). By contrast, if a variable is important to the classification, the prediction accuracy will drop after such a permutation, which we record as the MDA. Thus, the random forest analysis provides an "importance" rank ordering of biochemicals; we typically output the top 30 biochemicals in the list as potentially worthy of further investigation.

##### 11. Hierarchical Clustering

Hierarchical clustering is an unsupervised method for clustering the data, and can show large-scale differences. There are several types of hierarchical clustering and many distance metrics that can be used. A common method is complete clustering using the Euclidean distance, where each sample is a vector with all of the metabolite values. The differences seen in the cluster may be unrelated to the treatment groups or study design.

**12. Principal Components Analysis (PCA)**

Principal components analysis is an unsupervised analysis that reduces the dimension of the data. Each principal component is a linear combination of every metabolite and the principal components are uncorrelated. The number of principal components is equal to the number of observations.

The first principal component is computed by determining the coefficients of the metabolites that maximizes the variance of the linear combination. The second component finds the coefficients that maximize the variance with the condition that the second component is orthogonal to the first. The third component is orthogonal to the first two components and so on. The total variance is defined as the sum of the variances of the predicted values of each component (the variance is the square of the standard deviation), and for each component, the proportion of the total variance is computed. For example, if the standard deviation of the predicted values of the first principal component is 0.4 and the total variance = 1, then 100*0.4*0.4/1 = 16% of the total variance is explained by the first component. Since this is an unsupervised method, the main components may be unrelated to the treatment groups, and the "separation" does not give an estimate of the true predictive ability.

**13. Z-scores**

An intensity measurement for a metabolite by itself does not tell much. If for example a patient contains a blood glucose level of 300, this could be very good news if most people have blood glucose levels around 300, but less so if most people have levels around 100. In other words a measurement is meaningful only relative to the means of the sample or the population. This can be achieved by transforming the measurements into Z-scores which are expressed as standard deviations from the mean.

The Z-score, also called the standard score or normal score, is a dimensionless quantity derived by subtracting the control population mean from an individual raw score and then dividing the difference by the control population standard deviation. The Z-score indicates how many standard deviations an observation is above or below the mean of the control group. The Z-score is negative when the raw score is below the mean, positive when above. Since knowing the true mean and standard deviation of a control population is often unrealistic, the mean and standard deviation of the control population may be estimated using a random control sample.

Z-score =

where: x is a raw score to be standardized, μ is the mean of the control population, σ is the standard deviation of the control population

Subtracting the mean *centers* the distribution, and dividing by the standard deviation *standardizes* the distribution. The interesting properties of Z-scores are that they have a zero mean (effect of "centering") and a variance and standard deviation of 1 (effect of "standardizing"). This is because all distributions expressed in Z-scores have the same mean (0) and the same variance (1), so we can use Z-scores to compare observations coming from different distributions. When a distribution is normal most of the Z-scores (more than 99%) lay between the values of -3 and +3.

### mList+$^{TM}$ FINAL REPORT

#### Biochemical profiling of BPD plasma samples

##### UCAM-0312E-15MLBL+

CLIENT: Willem Ouwehand PhD

AUTHOR: Gregory Michelotti, PhD

APPROVAL: Bryan Wittmann, PhD

DATE: August 2, 2016

Metabolon, Inc. • 617 Davis Drive, Suite 400, Durham, NC 27713 • (919) 572-1711 www.metabolon.com • Contact:

Table of Contents

**Objective**

**Experimental Procedures**

**Results and Biological Interpretation**

*[Biological Interpretation is available as a component of mView service deliverable.]*

***Metabolite Summary and Significantly Altered Biochemicals***

**Study Parameters**

***Data Quality: Instrument and Process Variability***

**Appendix**

***Metabolon Platform***

***Statistical Methods and Terminology***

### Objective

#### *Purpose of Experiment*

The goal of the study was to identify metabolic differences in rare disease group samples that are associated with BPD samples from the Ouwehand group.

### Experimental Procedures

#### *Experimental design*

Global biochemical profiles were determined in human plasma samples associated with samples below.

| Matrix | Sample n | Description |
|---|---|---|
| Human Plasma | 58 | Willem Ouwehand's BPD samples |

### Results and Biological Interpretation

*[Biological Interpretation is available as a component of mView service deliverable.]*

#### *Metabolite Summary and Significantly Altered Biochemicals*

Numbers of compounds of known structural identity (named biochemicals) as well as compounds of unknown structural identity (unnamed biochemicals) detected in the present datasets are presented in the table below. Information related to biochemical pathways, chemical properties, and public databases are included in the electronic data files that accompany this report.

| *Matrix* | *Named Biochemicals* | *Unnamed Biochemicals* | *Total Biochemicals* |
|---|---|---|---|
| Human Plasma | 947 | 433 | 1380 |

### Study Parameters

#### *Data Quality: Instrument and Process Variability*

| *QC Sample* | *Measurement* | *Median RSD* |
|---|---|---|
| Internal Standards | Instrument Variability | 4% |
| Endogenous Biochemicals | Total Process Variability | 9% |

Instrument variability was determined by calculating the median relative standard deviation (RSD) for the internal standards that were added to each sample prior to injection into the mass spectrometers. Overall process variability was determined by calculating the median RSD for all endogenous metabolites (i.e., non-instrument standards) present in 100% of the Client Matrix samples, which are technical replicates of pooled client samples. Values for instrument and process variability as shown in the table above meet Metabolon's acceptance criteria.

### Appendix

#### *Metabolon Platform*

**Sample Accessioning:** Following receipt, samples were inventoried and immediately stored at -80$^{o}$C. Each sample received was accessioned into the Metabolon LIMS system and was assigned by the LIMS a unique identifier that was associated with the original source identifier only. This identifier was used to track all sample handling, tasks, results, etc. The samples (and all derived aliquots) were tracked by the LIMS system. All portions of any sample were automatically assigned their own unique identifiers by the LIMS when a new task was created; the relationship of these samples was also tracked. All samples were maintained at -80$^{o}$C until processed.

**Sample Preparation:** Samples were prepared using the automated MicroLab STAR® system from Hamilton Company. Several recovery standards were added prior to the first step in the extraction process for QC purposes. To remove protein, dissociate small molecules bound to protein or trapped in the precipitated protein matrix, and to recover chemically diverse metabolites, proteins were precipitated with methanol under vigorous shaking for 2 min (Glen Mills GenoGrinder 2000) followed by centrifugation. The resulting extract was divided into five fractions: two for analysis by two separate reverse phase (RP)/UPLC-MS/MS methods with positive ion mode electrospray ionization (ESI), one for analysis by

RP/UPLC-MS/MS with negative ion mode ESI, one for analysis by HILIC/UPLC-MS/MS with negative ion mode ESI, and one sample was reserved for backup. Samples were placed briefly on a TurboVap® (Zymark) to remove the organic solvent. The sample extracts were stored overnight under nitrogen before preparation for analysis.

**QA/QC:** Several types of controls were analyzed in concert with the experimental samples: a pooled matrix sample generated by taking a small volume of each experimental sample (or alternatively, use of a pool of well-characterized human plasma) served as a technical replicate throughout the data set; extracted water samples served as process blanks; and a cocktail of QC standards that were carefully chosen not to interfere with the measurement of endogenous compounds were spiked into every analyzed sample, allowed instrument performance monitoring and aided chromatographic alignment. Tables 1 and 2 describe these QC samples and standards. Instrument variability was determined by calculating the median relative standard deviation (RSD) for the standards that were added to each sample prior to injection into the mass spectrometers. Overall process variability was determined by calculating the median RSD for all endogenous metabolites (i.e., non-instrument standards) present in 100% of the pooled matrix samples. Experimental samples were randomized across the platform run with QC samples spaced evenly among the injections, as outlined in Figure 1.

**Table 1: Description of Metabolon QC Samples**

| Type | Description | Purpose |
|---|---|---|
| MTRX | Large pool of human plasma maintained by Metabolon that has been characterized extensively. | Assure that all aspects of the Metabolon process are operating within specifications. |
| CMTRX | Pool created by taking a small aliquot from every customer sample. | Assess the effect of a non-plasma matrix on the Metabolon process and distinguish biological variability from process variability. |
| PRCS | Aliquot of ultra-pure water | Process Blank used to assess the contribution to compound signals from the process. |
| SOLV | Aliquot of solvents used in extraction. | Solvent Blank used to segregate contamination sources in the extraction. |

**Table 2: Metabolon QC Standards**

| Type | Description | Purpose |
| --- | --- | --- |
| RS | Recovery Standard | Assess variability and verify performance of extraction and instrumentation. |
| IS | Internal Standard | Assess variability and performance of instrument. |

**Figure 1. Preparation of client-specific technical replicates.** A small aliquot of each client sample (colored cylinders) is pooled to create a CMTRX technical replicate sample (multi-colored cylinder), which is then injected periodically throughout the platform run. Variability among consistently detected biochemicals can be used to calculate an estimate of overall process and platform variability.

**Ultrahigh Performance Liquid Chromatography-Tandem Mass Spectroscopy (UPLC-MS/MS):** All methods utilized a Waters ACQUITY ultra-performance liquid chromatography (UPLC) and a Thermo Scientific Q-Exactive high resolution/accurate mass spectrometer interfaced with a heated electrospray ionization (HESI-II) source and Orbitrap mass analyzer operated at 35,000 mass resolution. The sample extract was dried then reconstituted in solvents compatible to each of the four methods. Each reconstitution solvent contained a series of standards at fixed concentrations to ensure injection and chromatographic consistency. One aliquot was analyzed using acidic positive ion conditions, chromatographically optimized for more hydrophilic compounds. In this method, the extract was gradient eluted from a C18 column (Waters UPLC BEH C18-2.1x100 mm, 1.7 µm) using water and methanol, containing 0.05% perfluoropentanoic acid (PFPA) and 0.1% formic acid (FA). Another aliquot was also analyzed using acidic positive ion conditions, however it was chromatographically optimized for more hydrophobic compounds. In this method, the extract was gradient eluted from the same afore mentioned C18 column using methanol, acetonitrile, water, 0.05% PFPA and 0.01% FA and was operated at an overall higher organic content. Another aliquot was analyzed using basic negative ion optimized conditions using a separate dedicated C18 column. The basic extracts were gradient eluted from the column using methanol and water, however with 6.5mM Ammonium Bicarbonate at pH 8. The fourth aliquot was analyzed via negative ionization following elution from a HILIC column (Waters UPLC BEH Amide 2.1x150 mm, 1.7 µm) using a gradient consisting of water and acetonitrile with 10mM Ammonium Formate, pH 10.8. The MS analysis alternated between MS and data-dependent $MS^n$ scans using dynamic exclusion. The scan range varied slighted between methods but covered 70-1000 m/z. Raw data files are archived and extracted as described below.

**Bioinformatics:** The informatics system consisted of four major components, the Laboratory Information Management System (LIMS), the data extraction and peak-identification software, data processing tools for QC and compound identification, and a collection of information interpretation and visualization tools for use by data analysts. The hardware and software foundations for these informatics components were the LAN backbone, and a database server running Oracle 10.2.0.1 Enterprise Edition.

**LIMS:** The purpose of the Metabolon LIMS system was to enable fully auditable laboratory automation through a secure, easy to use, and highly specialized system. The scope of the Metabolon LIMS system encompasses sample accessioning, sample preparation and instrumental analysis and reporting and advanced data analysis. All of the subsequent software systems are grounded in the LIMS data structures. It has been modified to leverage and interface with the in-house information extraction and data visualization systems, as well as third party instrumentation and data analysis software.

**Data Extraction and Compound Identification:** Raw data was extracted, peak-identified and QC processed using Metabolon's hardware and software. These systems are built on a web-service platform utilizing Microsoft's .NET technologies, which run on high-performance application servers and fiber-channel storage arrays in clusters to provide active failover and load-balancing. Compounds were identified by comparison to library entries of purified standards or recurrent unknown entities. Metabolon maintains a library based on authenticated standards that contains the retention time/index (RI), mass to charge ratio (*m/z)*, and chromatographic data (including MS/MS spectral data) on all molecules present in the library. Furthermore, biochemical identifications are based on three criteria: retention index within a narrow RI window of the proposed identification, accurate mass match to the library +/- 10 ppm, and the MS/MS forward and reverse scores between the experimental data and authentic standards. The MS/MS scores are based on a comparison of the ions present in the experimental spectrum to the ions present in the library spectrum. While there may be similarities between these molecules based on one of these factors, the use of all three data points can be utilized to distinguish and differentiate biochemicals. More than 3300 commercially available purified standard compounds have been acquired and registered into LIMS for analysis on all platforms for determination of their analytical characteristics. Additional mass spectral entries have been created for structurally unnamed biochemicals, which have been identified by virtue of their recurrent nature (both chromatographic and mass spectral). These compounds have the potential to be identified by future acquisition of a matching purified standard or by classical structural analysis.

**Curation:** A variety of curation procedures were carried out to ensure that a high quality data set was made available for statistical analysis and data interpretation. The QC and curation processes were designed to ensure accurate and consistent identification of true chemical entities, and to remove those representing system artifacts, mis-assignments, and background noise. Metabolon data analysts use proprietary visualization and interpretation software to confirm the consistency of peak identification among the various samples. Library matches for each compound were checked for each sample and corrected if necessary.

**Metabolite Quantification and Data Normalization:** Peaks were quantified using area-under-the-curve. For studies spanning multiple days, a data normalization step was performed to correct variation resulting from instrument inter-day tuning differences. Essentially, each compound was corrected in run-day blocks by registering the medians to equal one (1.00) and normalizing each data point proportionately (termed the "block correction"; Figure 2). For studies that did not require more than one day of analysis, no normalization is necessary, other than for purposes of data visualization. In certain instances, biochemical data may have been normalized to an additional factor (e.g., cell counts, total protein as determined by Bradford assay, osmolality, etc.) to account for differences in metabolite levels due to differences in the amount of material present in each sample.

**Figure 2: Visualization of data normalization steps for a multiday platform run.**

#### *Statistical Methods and Terminology*

**Statistical Calculations:** For many studies, two types of statistical analysis are usually performed: (1) significance tests and (2) classification analysis. Standard statistical analyses are performed in ArrayStudio on log transformed data. For those analyses not standard in ArrayStudio, the programs R (http://cran.r-project.org/) or JMP are used. Below are examples of frequently employed significance tests and classification methods followed by a discussion of p- and q-value significance thresholds.

#### 1. Welch's two-sample *t*-test

Welch's two-sample *t*-test is used to test whether two unknown means are different from two independent populations.

This version of the two-sample *t*-test allows for unequal variances (variance is the square of the standard deviation) and has an *approximate t*-distribution with degrees of freedom estimated using Satterthwaite's approximation. The test statistic is given by *t* , and the degrees of freedom is given by , where , are the sample means, $s_1$, $s_2$, are the sample standard deviations, and $n_1$, $n_2$ are the samples sizes from groups 1 and 2, respectively. We typically use a two-sided test (tests whether the means are different) as opposed to a one-sided test (tests whether one mean is greater than the other).

#### 2. Matched Pairs *t*-test

The matched pairs *t*-test is used to test whether two unknown means are different from paired observations taken on the same subjects.

The matched pairs *t*-test is equivalent to the one-sample *t*-test performed on the differences of the observations taken on each subject (i.e., calculate ($x_1 - x_2$) for each subject; test whether the mean difference is zero or not). The test statistic is given by , with $n - 1$ degrees of freedom, where , are the sample means for groups 1 and 2, respectively, $s_d$ is the standard deviation of the differences, *n* is the number of *subjects* (so there are 2*n* observations).

#### 3. One-way ANOVA

ANOVA stands for analysis of variance. For ANOVA, it is assumed that all populations have the same variances. One-way ANOVA is used to test whether at least two unknown means are all equal or whether at least one pair of means is different. For the case of two means, ANOVA gives the same result as a two-sided *t*-test with a pooled estimate of the variance.

An ANOVA uses an F-test which has two parameters – the numerator degrees of freedom and the denominator degrees of freedom. The degrees of freedom in the numerator are equal to $g - 1$, where *g* is the number of groups. If *n* is the total number of observations ($n_1 + n_2$), then, the denominator degrees of freedom is equal to $n - g$. The F-statistic is the ratio of the between-groups variance to the within-groups variance, hence the higher the F-statistic the more evidence we have that the means are different.

Often within ANOVA, one performs linear contrasts for specific comparisons of interest. For example, suppose we have three groups A, B, C, then examples of some contrasts are A vs. B, the average of A and B vs. C, etc. For single-degree of freedom contrasts, these give the same result as a two-sided *t*-test with the pooled estimate of the variance from the ANOVA and degrees of freedom $n - g$. Below, we show the three formulas for A vs. B from a three group design as shown above. The numerator is same in each case, but the denominator differs by the estimates of the variances, and the degrees of freedom are different for each (if the theoretical assumptions hold, then the contrast has the most power, as it has the largest degrees of freedom).

Welch's two-sample *t*-test
By $t$ , and the degrees of freedom is given by

Two-sample *t*-test with pooled estimate of variance from A and B
where , where the degrees of freedom is $n_A + n_B - 2$.
The contrast from the ANOVA,
where , where the degrees of freedom is given by where the degrees of freedom is $n_A + n_B + n_C - 3$.

**4. Two-way ANOVA**

ANOVA stands for analysis of variance. For ANOVA, it is assumed that all populations have the same variances. For a two-way ANOVA, three statistical tests are typically performed: the main effect of each factor and the interaction. Suppose we have two factors A and B, where A represent the genotype and B represent the diet in a mouse study. Suppose each of these factors has two levels (A: wild type, knock out; B: standard diet, high fat diet). For this example, there are 4 combinations ("treatments"): A1B1, A1B2, A2B1, A2B2. The overall ANOVA F-test gives the p-value for testing whether all four of these means are equal or whether at least one pair is different. However, we are also interested in the effect of the genotype and diet. A main effect is a contrast that tests one factor across the levels of the other factor. Hence the A main effect compares (A1B1 + A1B2)/2 vs. (A2B1 + A2B2)/2, and the B-main effect compares (A1B1 + A2B2)/2 vs. (A1B2 + A2B2)/2. The interaction is a contrast that tests whether the mean difference for one factor depends on the level of the other factor, which is (A1B2 + A2B1)/2 vs. (A1B1 + A2B2)/2.

Some sample plots follow. For the first plot, there is a B main effect, but no A main effect and no interaction, as the effect of B does not depend on the level of A. For the second plot, notice how the mean difference for B is the same at each level of A and the difference in A is the same for each level of B, hence there is no statistical interaction. The final plot also has main effects for A and B, but here also has an interaction: we see the effect of B depends on the level of A (0 for A1 but 2 for A2), i.e., the effect of the diet depends on the genotype. We also see here the interpretation of the main effects depends on whether there is an interaction or not.

**5. Two-way Repeated Measures ANOVA**

This is typically an ANOVA where one factor is applied to each subject and the second factor is a time point. See two-way ANOVA as many of the details are similar except that the model takes into account the repeated measures, i.e., the treatments are given to the same subject over time. The two main effects and the interaction are assessed, with particular interest to the interaction, as this shows where the time profiles are parallel or not for the treatments (parallel mean no interaction).

One additional note, the standard analysis assumes a condition referred to as compound symmetry, which assumes the correlation between each pair of levels of the repeated-measures factor is the same. Thus, for the case of time, it assumes the correlation is the same between time points 1 and 2, 1 and 3, and 2 and 3.

**6. Correlation**

Correlation measures the strength and direction of a *linear* association between two variables. The statistical test for correlation tests whether the true correlation is zero or not.

The square of the correlation is the percentage of the total variation explained by a linear relationship between the two variables. Thus, with large sample sizes there may be a sample correlation of 0.1 that is statistically significant. This means we have high confidence that the true correlation is zero, however, only 100*(0.1*0.1)% = 1% of the variation of one variable is explained by a linear relationship with the other variable, so while there is an association, it has little predictive ability.

**7. Hotelling's $T^2$ test**

The Hotelling's $T^2$ test is a multivariate generalization of the *t*-test, but here we are testing whether the mean vectors are different or not (the vector consists of multiple metabolites).

The Hotelling statistic is: , where $n_x$ and $n_y$ are the numbers of samples in each group, is the mean vector of the variables from group 1, is the mean vector of variables from group 2 and **S** is the pooled estimate of the variance-covariance matrix of the variables. This analysis assumes the underlying variance-covariance matrix is the same for each group. Notice that in the case of uncorrelated variables, this is simply a weighted average of the squared mean differences with weights inversely proportional to the sample variances (i.e., the metabolites less variable within a group are given higher weights).

#### 8. p-values

For statistical significance testing, p-values are given. The lower the p-value, the more evidence we have that the null hypothesis (typically that two population means are equal) is not true. If "statistical significance" is declared for p-values less than 0.05, then 5% of the time we incorrectly conclude the means are different, when actually they are the same.

The p-value is the probability that the test statistic is at least as extreme as observed in this experiment given that the null hypothesis is true. Hence, the more extreme the statistic, the lower the p-value and the more evidence the data gives against the null hypothesis.

#### 9. q-values

The level of 0.05 is the false positive rate when there is one test. However, for a large number of tests we need to account for false positives. There are different methods to correct for multiple testing. The oldest methods are family-wise error rate adjustments (Bonferroni, Tukey, etc.), but these tend to be extremely conservative for a very large number of tests. With gene arrays, using the False Discovery Rate (FDR) is more common. The family-wise error rate adjustments give one a high degree of confidence that there are zero false discoveries. However, with FDR methods, one can allow for a small number of false discoveries. The FDR for a given set of compounds can be estimated using the q-value (see Storey J and Tibshirani R. (2003) Statistical significance for genomewide studies. Proc. Natl. Acad. Sci. USA 100: 9440-9445; PMID: 12883005).

In order to interpret the q-value, the data must first be sorted by the p-value then choose the cutoff for significance (typically p<0.05). The q-value gives the false discovery rate for the selected list (i.e., an estimate of the proportion of false discoveries for the list of compounds whose p-value is below the cutoff for significance). For Table 1 below, if the whole list is declared significant, then the false discovery rate is approximately 10%. If everything from Compound 079 and above is declared significant, then the false discovery rate is approximately 2.5%.
Table 1: Example of q-value interpretation

#### 10. Random Forest

Random forest is a supervised classification technique based on an ensemble of decision trees (see Breiman L. (2001) Random Forests. Machine Learning. 45: 5-32; http://link.springer.com/article/10.1023%2FA%3A1010933404324). For a given decision tree, a random subset of the data with identifying true class information is selected to build the tree ("bootstrap sample" or "training set"), and then the remaining data, the "out-of-bag" (OOB) variables, are passed down the tree to obtain a class prediction for each sample. This process is repeated thousands of times to produce the forest. The final classification of each sample is determined by computing the class prediction frequency ("votes") for the OOB variables over the whole forest. For example, suppose the random forest consists of 50,000 trees and that 25,000 trees had a prediction for sample 1. Of these 25,000, suppose 15,000 trees classified the sample as belonging to Group A and the remaining 10,000 classified it as belonging to Group B. Then the votes are 0.6 for Group A and 0.4 for Group B, and hence the final classification is Group A. This method is unbiased since the prediction for each sample is based on trees built from a subset of samples that do not include that sample. When the full forest is grown, the class predictions are compared to the true classes, generating the "OOB error rate" as a measure of prediction accuracy. Thus, the prediction accuracy is an unbiased estimate of how well one can predict sample class in a new data set. Random forest has several advantages – it makes no parametric assumptions, variable selection is not needed, it does not overfit, it is invariant to transformation, and it is fairly easy to implement with R.

To determine which variables (biochemicals) make the largest contribution to the classification, a "variable importance" measure is computed. We use the "Mean Decrease Accuracy" (MDA) as this metric. The MDA is determined by randomly permuting a variable, running the observed values through the trees, and then reassessing the prediction accuracy. If a variable is not important, then this procedure will have little change in the accuracy of the class prediction (permuting random noise will give random noise). By contrast, if a variable is important to the classification, the prediction accuracy will drop after such a permutation, which we record as the MDA. Thus, the random forest analysis provides an "importance" rank ordering of biochemicals; we typically output the top 30 biochemicals in the list as potentially worthy of further investigation.

#### 11. Hierarchical Clustering

Hierarchical clustering is an unsupervised method for clustering the data, and can show large-scale differences. There are several types of hierarchical clustering and many distance metrics that can be used. A common method is complete clustering using the Euclidean distance, where each sample is a vector with all of the metabolite values. The differences seen in the cluster may be unrelated to the treatment groups or study design.

#### 12. Principal Components Analysis (PCA)

Principal components analysis is an unsupervised analysis that reduces the dimension of the data. Each principal component is a linear combination of every metabolite and the principal components are uncorrelated. The number of principal components is equal to the number of observations.

The first principal component is computed by determining the coefficients of the metabolites that maximizes the variance of the linear combination. The second component finds the coefficients that maximize the variance with the condition that the second component is orthogonal to the first. The third component is orthogonal to the first two components and so on. The total variance is defined as the sum of the variances of the predicted values of each component (the variance is the square of the standard deviation), and for each component, the proportion of the total variance is computed. For example, if the standard deviation of the predicted values of the first principal component is 0.4 and the total variance = 1, then 100*0.4*0.4/1 = 16% of the total variance is explained by the first component. Since this is an unsupervised method, the main components may be unrelated to the treatment groups, and the “separation” does not give an estimate of the true predictive ability.

#### 13. Z-scores

An intensity measurement for a metabolite by itself does not tell much. If for example a patient contains a blood glucose level of 300, this could be very good news if most people have blood glucose levels around 300, but less so if most people have levels around 100. In other words a measurement is meaningful only relative to the means of the sample or the population. This can be achieved by transforming the measurements into Z-scores which are expressed as standard deviations from the mean.

The Z-score, also called the standard score or normal score, is a dimensionless quantity derived by subtracting the control population mean from an individual raw score and then dividing the difference by the control population standard deviation. The Z-score indicates how many standard deviations an observation is above or below the mean of the control group. The Z-score is negative when the raw score is below the mean, positive when above. Since knowing the true mean and standard deviation of a control population is often unrealistic, the mean and standard deviation of the control population may be estimated using a random control sample.

Z-score =
where: x is a raw score to be standardized, μ is the mean of the control population, σ is the standard deviation of the control population

Subtracting the mean *centers* the distribution, and dividing by the standard deviation *standardizes* the distribution. The interesting properties of Z-scores are that they have a zero mean (effect of "centering") and a variance and standard deviation of 1 (effect of "standardizing"). This is because all distributions expressed in Z-scores have the same mean (0) and the same variance (1), so we can use Z-scores to compare observations coming from different distributions. When a distribution is normal most of the Z-scores (more than 99%) lay between the values of -3 and +3.
